# Germinal center B cells separate signals for selection and division like compact discs

**DOI:** 10.1101/2020.01.29.925479

**Authors:** Michael Meyer-Hermann

## Abstract

Selection of B cells in germinal center (GC) reactions is pivotal to the generation of high affinity antibodies and memory B cells. Knowledge about B cell selection is limited to individual interactions and lacks global understanding. Here, seemingly contradictory experiments are unified in a common concept by separation of signals for the frequency of fate decision from the strength of cell division, which can then be controlled independently, similar to the separation of frequency and amplitude in digital audio signals. Based on this concept of information processing, three theories of B cell selection are developed that can be distinguish by predicted experiments drawn from computer simulations. Understanding encoding of information in molecular states is critical for targeted immune interventions.

## Introduction

Germinal centers (GC) are specialized environments in lymphoid tissues that give rise to high affinity antibodies (Victora *et al.*, 2012). The creation of new antibodies that previously did not exist in an organism was well described in many details including GC morphology (Camacho *et al.*, 1998) with dark and light zone (DZ and LZ), inter-zonal cell motility (Allen *et al.*, 2007; Schwickert *et al.*, 2007; Hauser *et al.*, 2007; Figge *et al.*, 2008), B cell (BC) mutation (MacLennan, 1994; Oprea *et al.*, 2000), T follicular helper cell (Tfh)-dependent selection and division (Victora *et al.*, 2010; Meyer-Hermann *et al.*, 2012), and clonal diversity (Tas *et al.*, 2016). The widely accepted picture is that BCs undergo somatic hypermutation in the GC-DZ, where BC division is dominant, while BC selection is concentrated in the GC-LZ, where native antigen is presented on follicular dendritic cells (FDC) and cognate interactions with Tfh cells take place.

BC selection is based on two steps (Lindhout *et al.*, 1995): at first, BCs have to collect antigen from FDCs. Secondly, BCs have to get selection signals from Tfh. We proposed that Tfh are able to sense the affinity of the BC receptor (BCR) by the density of antigen-derived peptide-loaded MHC (pMHC) on the BC surface (Meyer-Hermann *et al.*, 2006): higher affinity BCs collect antigen from FDC more efficiently, subsequently process more antigen, and finally present higher numbers of pMHC on their surface. Thus, the density of pMHC reflects the BCR affinity for the antigen. Tfhs can interact with many competing pMHC-presenting BCs and polarize to the BC with highest pMHC density (Depoil *et al.*, 2005), respective highest affinity. When the BC received a critical amount of Tfh signals it gets positively selected. BCs failing to do so activate apoptosis. This theory of BC selection based on competition for Tfh help was supported by experimental findings (Allen *et al.*, 2007; Victora *et al.*, 2010).

While this theory of BC selection is state-of-the-art, the actual signaling of BCs in the course of BC mutation and subsequent selection remains elusive. Only recently, BC signaling came into the focus of research and pointed to a critical role of molecules like c-Myc (Calado *et al.*, 2012; Heinzel *et al.*, 2017; Luo *et al.*, 2018), FoxO1 (Sander *et al.*, 2015; Inoue *et al.*, 2017), and mTOR (Ersching *et al.*, 2017). All these factors are involved in BC signaling downstream of BCR and CD40 ligation by Tfhs, which is critical for BC survival. The relative contribution of BCR and Tfh signals to the regulation of these factors was disentangled and a synergy of both pathways was identified (Luo *et al.*, 2018; Turner *et al.*, 2018). In addition, a positive feedback loop between CD40 ligation and ICOSL upregulation was described (Liu *et al.*, 2015) giving rise to a non-linear relationship of BC stimulation and downstream signaling and makes interpretation of experimental results less intuitive.

One constraint that determines the architecture of signal progression is the observation of a differential number of divisions (DND). Based on *in vivo* observations (Victora *et al.*, 2010) and computer simulation (Meyer-Hermann *et al.*, 2012), it was found that the interaction between Tfh and BCs gives rise to not only BC selection and subsequent differentiation, but that also the number of subsequent divisions, rather than being a random number or an endogenous BC property, is controlled by these very same interactions with Tfhs and ranges between one and six divisions. While, this was confirmed in specific experiments (Gitlin *et al.*, 2014; Finkin *et al.*, 2019), it also bears a structural and unresolved limitation: assuming selection was determined by a particular signal *X*, which is integrated from BC antigen uptake and/or multiple interactions with different Tfh, a threshold signal *X*_th_ would exist at which BC selection and fate decision is initiated. Then, all selected BCs would exhibit a similar signal level *X* ≈ *X*_th_ at the moment of selection. If this very same signal was also responsible for the induced DND in the DZ, all selected BCs would divide an equal amount of times, which is not the case (Gitlin *et al.*, 2014). Thus, this selection signal *X* cannot encode the subsequent DND, the information about which must be encoded in a different signaling molecule *Y*, which either dynamically controls the threshold *X*_th_(*Y*) or directly controls DND.

The relative role of BCR and Tfh signals was disentangled by providing antigen to BCs in a BCR-independent way taking advantage of the DEC205-receptor (Victora *et al.*, 2010). This receptor is expressed on BCs and allows for antigen-uptake irrespective of interactions of BCRs with antigen hold on FDCs. BCs that take up antigen via the DEC205-receptor exhibited high antigen-pMHC presentation, prolonged LZ passage times, and a division burst in the DZ. While the division burst can be understood by increased Tfh help to high pMHC-presenting BCs, the prolonged LZ passage time was never understood and actually contradicts the general expectation that high affinity BCs would get selected faster than low affinity BC. One might consider this experimental system artificial and non-physiological. However, a realistic theory of BC selection has to properly describe GCs permissive of low affinity BCs (Kuraoka *et al.*, 2016; Silver *et al.*, 2018) still allowing for efficient affinity maturation, diversity of GC clonal dominance (Tas *et al.*, 2016) still allowing for clonal bursts, Tfh signal-dependent DND upon selection (Gitlin *et al.*, 2014) without losing the relevance of BCR signaling for selection, and also reflect extreme stimulation settings like antigen-uptake via the DEC205-receptor (Victora *et al.*, 2010). A consistent theory, explaining these seemingly conflicting GC properties does not exist (Oprea & Perelson, 1997; Zhang & Shakhnovich, 2010; Meyer-Hermann *et al.*, 2012; Wang *et al.*, 2016; de Boer & Perelson, 2017; Meyer-Hermann, 2019).

Here, theories of BC information processing in agreement with all aforementioned experimental results are proposed. The long-standing lack of mechanistic understanding of GC selection is resolved based on a concept of information processing that separates the signals for the frequency of selection from the strength of subsequent division. The theories are tested for observable implications in computer simulations and experimental tests are proposed to determine which version of signal separation was realized in nature. The structural necessity of signal separation has implications for experiments aiming at manipulation of GC reactions for clinical purposes.

## Results

### Three theories separating selection and division signals

A spatio-temporal GC simulation framework was generated in which each BC, FDC, and Tfh is represented as an individual object (Meyer-Hermann *et al.*, 2012; Meyer-Hermann, 2014; Binder & Meyer-Hermann, 2016; Meyer-Hermann, 2019). In this framework, BCs divide and mutate in the DZ and interact with FDCs and Tfhs in the LZ. The subsequent BC fate, i.e. apoptosis or recycling to the DZ phenotype, is determined by signals collected from multiple FDCs and Tfhs. The signaling theories described below detail the mechanisms of BC selection (summarized in Figure 1) and were implemented for each cell object (see Supplementary Methods and Supplementary Table 1 for theory-specific parameters), inducing a diversity of cell behaviors depending on the individually experienced interactions. All presented theories have in common that they separate the signals for BC fate decision and the number of BC divisions.

**Figure 1:**
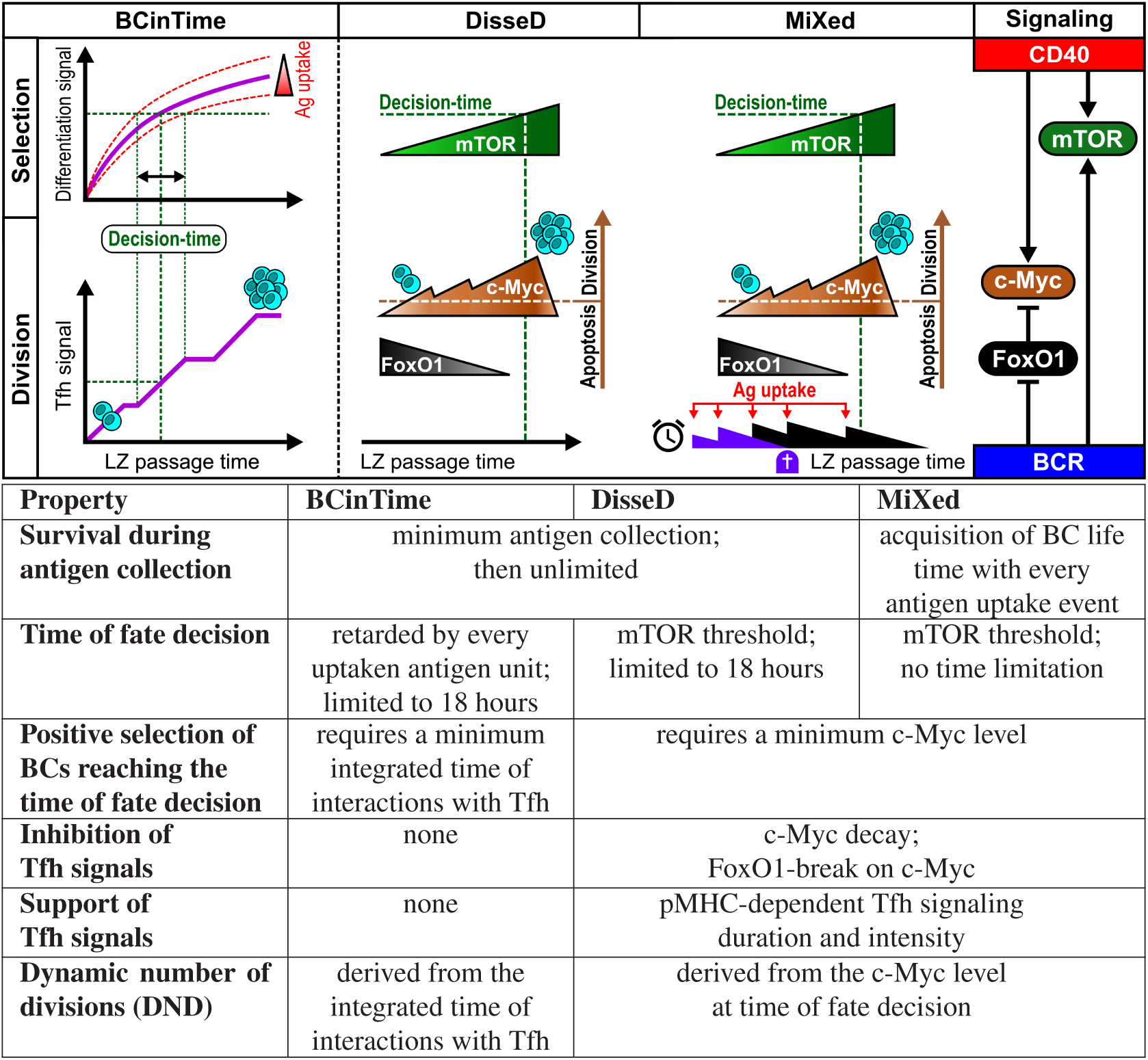
Scheme of the separation of selection and division signals in the three theories. During their journey through the LZ (*LZ passage time*), each simulated BC interacts with FDCs and Tfhs and collect signals, which determine the time of fate decision and their subsequent fate. In the BCinTime-theory, a non-specified differentiation signal is upregulated in dependence on the amount of uptaken antigen. This determines the time of fate decision. The amount of Tfh signals received by this time determines the subsequent number of divisions. In the DisseD- and the MiXed-theory the signals are made explicit and interact as depicted at the top right in each simulated BC: mTOR is upregulated in dependence on BCR and Tfh (CD40). cMyc is upregulated in dependence on Tfh (CD40) signals after BCR-signaling dependent release of the FoxO1-brake. Fate decision is induced when mTOR meets a threshold expression level. At this time, BCs with c-Myc, which is subject to degradation, above a threshold survive. The number of subsequent divisions is determined by the c-Myc level for surviving BCs. MiXeddiffers from DisseDby the additional requirement that BCs failing to take up antigen may die before mTOR reaches the fate decision threshold. Implementation in the three theories is made more concrete in the table. See Supplementary Table 1 for parameter values.

The DisseD-theory stands for Different signals for Selection and Division. As the name states, selection and DND are based on the signaling levels of two different factors, mTOR and c-Myc, respectively (Figure 1). In LZ-BCs, mTOR is upregulated in a BCR dependent manner (see Supplementary Eq. (11)). During Tfh-BC-interactions, BCR and Tfh signals synergize to upregulate mTOR more efficiently. When mTOR reaches a threshold expression level, BC fate decision is initiated. The achieved level of c-Myc determines the subsequent DND in the DZ (see Supplementary Eq. (12)). During the mTOR limited LZ passage time, c-Myc is upregulated in a Tfh-dependent manner (see Supplementary Eq. (10)), where its upregulation is inhibited by FoxO1. FoxO1 is a transcription factor associated with gene expression during BC development (Dengler *et al.*, 2008) and is dynamically regulated in GC reactions (Dominguez-Sola *et al.*, 2015) where FoxO1 is high in DZ-BCs and low in LZ-BCs (Sander *et al.*, 2015). Thus, c-Myc inhibition by FoxO1 was assumed high when BCs enter the LZ. The release of the FoxO1brake depends on BCR signals (see Supplementary Eq. (4)), such that c-Myc upregulation also relies on synergy of BCR and Tfh signals, but with different dynamics. The DisseD-theory is summarized in Figure 1 and translated into a set of differential equation solved for each cell object.

The BCinTime-theory is based on a rather simple assumption: The LZ passage time is determined by the efficiency of antigen-uptake from FDCs. With every piece of antigen that is taken up, the respective BC gets additional time until the fate decision is taken. This is realized by a phenomenological signal upregulated in LZ-BCs with a rate depending on the amount of uptaken antigen (see Supplementary Eq. (8)). With every additional antigen, the rate of signal upregulation is reduced. When the signal reaches a threshold value, interactions with Tfhs or FDCs are stopped and the fate decision is induced. The DND in the DZ is determined by the amount of Tfh signals accumulated until the time of fate decision (see Supplementary Eqs. (9) and (12)).

In the MiXed-theory, the mechanisms described in Figure 1 are used as in the DisseD-theory, complemented by an element of the BCinTime-theory further limiting the LZ passage time: After acquisition of the LZ phenotype, BC have an initial finite life time, during which they have to manage to catch some antigen from FDCs. Failure to do so implies irreversible activation of apoptosis. With each successful antigen-uptake, the life time is prolonged by a fixed time period. Thus, in repeated cycles, each BC has to capture new antigen within its limited life time. Surviving BCs continue to do so until they reach the mTOR level for fate decision. As in the DisseD-theory, c-Myc levels acquired by that time determine the DND.

### Basic GC characteristics are similar in all three theories

At first, generally accepted characteristics of the GC reaction are tested in the three theories in order to establish confidence that the computer simulations are reflecting real GC reactions. Subsequently, qualitative and quantitative differences of the three theories will be highlighted (summarized in Table 1). The latter will help to design experiments validating and distinguishing the three theories.

**Table 1:** Experimental observations and how they are reflected in the three theories.

| Observation/Reference | BCinTime | DisseD | MiXed | Figure |
| --- | --- | --- | --- | --- |
| GC size (Liu *et al.*, 1991; Hollowood & Macartney, 1992) | Reasonable GC size and population kinetics |  |  | 2a |
| GC shutdown (Zhang *et al.*, 2013) | fast | slow | slow | 2a |
| Robustness against antigen concentration (Vora *et al.*, 1997; Hannum *et al.*, 2000) | low | high | high | 2b,e;<br>S4a-d |
| Affinity maturation (Victora *et al.*, 2010) | well reproduced |  |  | 2d |
| DZ to LZ ratio is 2 (Victora *et al.*, 2010) | well reproduced |  |  | 3a-c |
| Competition for Tfh (prediction) | low | high | high | 2e |
| Mean DND of 2 and variation between 1 and 6 (Gitlin *et al.*, 2014) | well reproduced |  |  | 2h,i;<br>3g; S2i |
| Asymmetry of DZ to LZ and LZ to DZ transmigration (Victora *et al.*, 2010) | well reproduced |  |  | 4a |
| LZ passage time of high affinity BCs (prediction) | longer | shorter | shorter | 4b |
| LZ passage time of deleted BCs (prediction) | shorter | longer | shorter | 4c |
| Number of Tfh contacts per LZ passage in deleted BCs (prediction) | less | more | less | 4d |
| LZ accumulation and DZ burst in anti-DEC205-OVA experiments (Victora *et al.*, 2010) | well reproduced |  |  | 3a-c |
| Stop of affinity maturation in anti-DEC205-OVA experiments (Victora *et al.*, 2010) | well reproduced |  |  | S3 |
| mTOR-overexpressing BCs have competitive disadvantage (Ersching *et al.*, 2017) | qualitatively reproduced |  |  | S4e,f |
| Clonal diversity (Tas *et al.*, 2016) | qualitatively reproduced |  |  | 5a-c |
| Existence of monoclonal and multiclonal GCs (Tas *et al.*, 2016) | yes, frequent monoclonal | yes | yes, frequent multiclonal | 5d,e |
| Clonal dominance day 17 of the GC reaction (prediction) | higher | lower | intermediate | 5f,g |

The GC peak size is similar in all three theories (Figure 2a). However, the GC population is reduced faster for BCinTime. As antigen on FDC gets consumed by BCs, access to antigen becomes more difficult (Figure 2b,c). The reduced amount of collected antigen has a double negative effect on BC survival: It reduces the LZ passage time, which increases the fraction of LZ-BCs dying by apoptosis. But even those BCs that survived get lower Tfh signal levels, such that they also divide less in the DZ. Thus, the faster reduction of the GC population is an inherent result of the BCinTime-theory.

**Figure 2:**
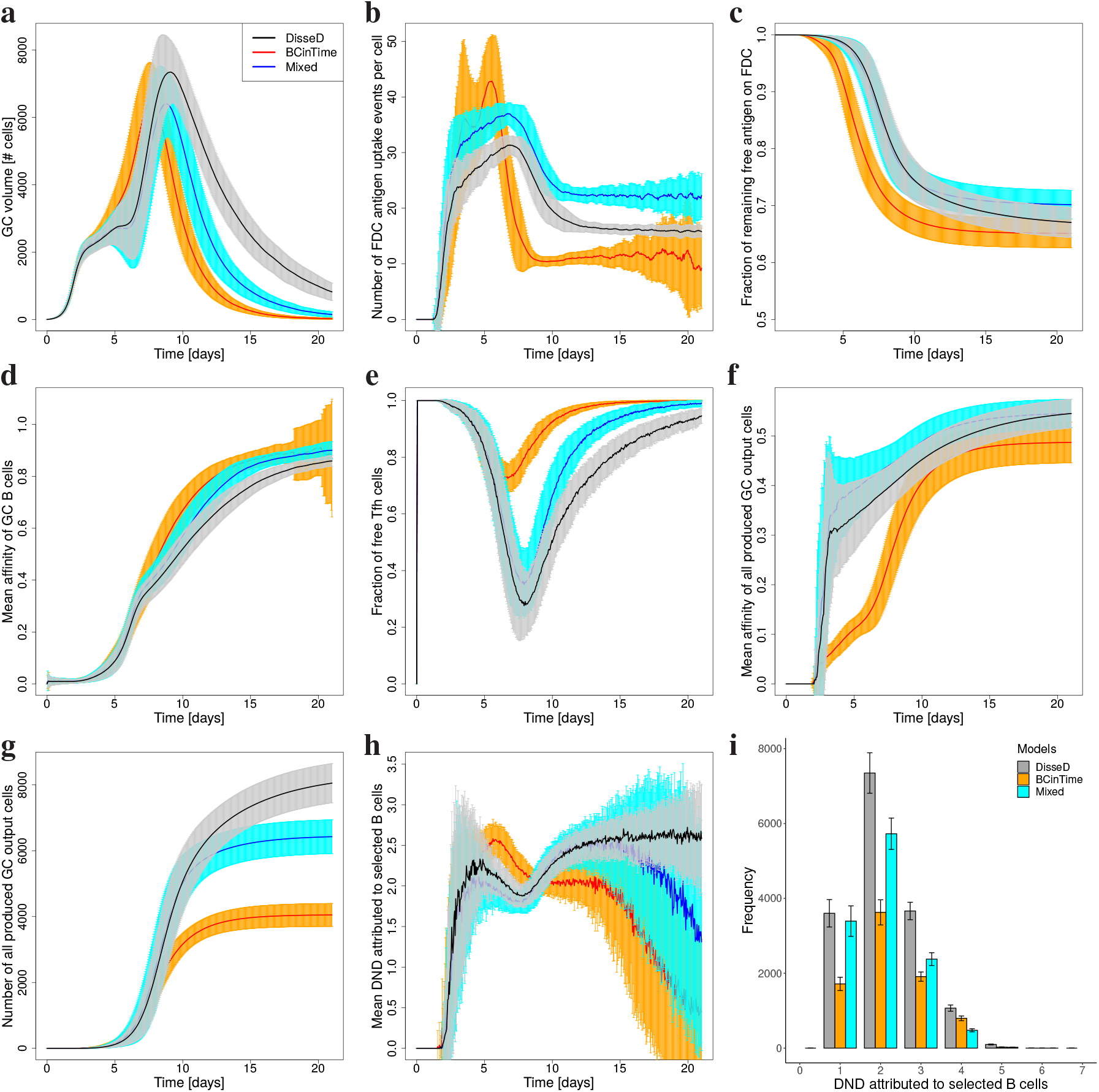
General GC characteristics of the three models. **(a)** Population kinetics, **(b)** antigen collection and **(c)** consumption, **(d)** mean affinity of GC-BCs, **(e)** competition for Tfhs, **(f)** mean affinity and **(g)** number of all output cells produced per GC, **(h)** mean and **(i)** distribution of the number of divisions attributed to selected BCs. DisseD-(black), BCinTime-(red), and MiXed-theory (blue). Mean and standard deviation over 100 GC simulations (a-g,i) and over all GC-BCs in a single representative simulation (h).

Despite this difference, affinity maturation of GC BCs is similar (Figure 2d). While competition for antigen is critical in the BCinTime-theory, this is equilibrated by less competition for Tfh help (Figure 2e). The lower affinity of output cells during the first days of the reaction (Figure 2f) points to a more permissive selection in the phase of abundant antigen. The GC tries to catch up when antigen availability gets limiting, as reflected in slightly accelerated affinity maturation (Figure 2d). However, the ultimate affinity of output cells is kept lower (Figure 2f), which is a result of the reduced total number of BCs passing through the GC, which infers less total output per GC in the late reaction (Figure 2g).

In DisseD and MiXed, the information on antigen taken up with the BCR is translated into the speed of release of the FoxO1-brake on c-Myc (Figure 1; Supplementary Figure 1a,b,f,g) and stored in the level of pMHC-presentation. The latter determines the frequency and the intensity of signals received by Tfh, which is reflected in the rate of mTOR and c-Myc upregulation (Supplementary Figure 1b,c,g,h). c-Myc levels at the time of fate decision, which is determined by mTOR reaching a threshold value, show a clear separation of deleted and selected BCs, where the separation is more pronounced in the MiXed-theory (Supplementary Figure 1d,e,i,j). The range of c-Myc levels in the subset of selected BCs encodes the DND.

The DND has a mean value around two with a distribution of values in the range of 1-6 (Figure 2h,i), both in agreement with (Gitlin *et al.*, 2014). Low affinity BCs divide less than high affinity BCs in all theories. While for DisseD and MiXed, DND drops in the phase of strong competition for Tfh help, in BCinTime, DND is increasing in the same phase and drops afterwards (Figure 2h). This reflects the critical role of antigen availability in the BCinTime-theory.

### anti-DEC205-OVA-experiments explained *in silico*

It is possible to provide antigen to BCs expressing the DEC205-receptor via injection of anti-DEC205-OVA. The BCs then internalize, process and present the antigen as pMHC on their surface just like they do with antigen taken up via the BCR but without activating the signaling machinery downstream of BCR and without the need to actually interact with FDCs to catch antigen. Such experiments *in vivo* disentangled the relative roles of Tfh and FDCs in BC selection (Victora *et al.*, 2010). anti-DEC205-OVA injection induced an accumulation of BCs in the LZ for 12 hours before a strong burst of BC division was induced in the DZ, as reflected in the dynamics of the DZ to LZ ratio (Victora *et al.*, 2010). Previous GC theories, like the LEDA model (Meyer-Hermann *et al.*, 2012), were able to explain the DZ burst but failed to explain the LZ accumulation without additional assumptions, such as increased interaction times with Tfh or equivalent, which were not supported by experimental data. The failure to reproduce LZ accumulation *in silico* was due to the lack of a detailed selection model. Next, consistency of the three selection theories with LZ accumulation and DZ burst were tested.

Without any further assumption, injection of anti-DEC205-OVA *in silico* leads to an accumulation of BCs in the LZ and to a subsequent proliferation burst in the DZ consistent with the experimental data in all theories (Figure 3a-c) but with subtle differences. LZ accumulation is robust against changes of the amount of DEC205-positive BCs in the simulation in the BCinTime-theory but not in the DisseD- and MiXed-theory, for which the LZ accumulation phase is extended and DZ expansion is reduced with more DEC205-positive BCs (Supplementary Figure 2a-c) because stronger competition for Tfhs leads to slower upregulation of mTOR and c-Myc. The DZ burst relies on dynamics in DND (Figure 3g; Supplementary Figure 2i) with lower peak DND in the BCinTime-theory. While in DisseD and BCinTime, the GC relaxes to normal within 3 days, the advantage of DEC205-positive BCs persists for longer times in the MiXed-theory (Figure 3c). Similar to the *in vivo* case, the DEC205-negative population exhibits only small modulations by the *in silico* injection. The fewer DEC205-positive BCs are in the system, the weaker the perturbation of the GC and the effect on DEC205-negative BCs (Supplementary Figure 2a-c).

**Figure 3:**
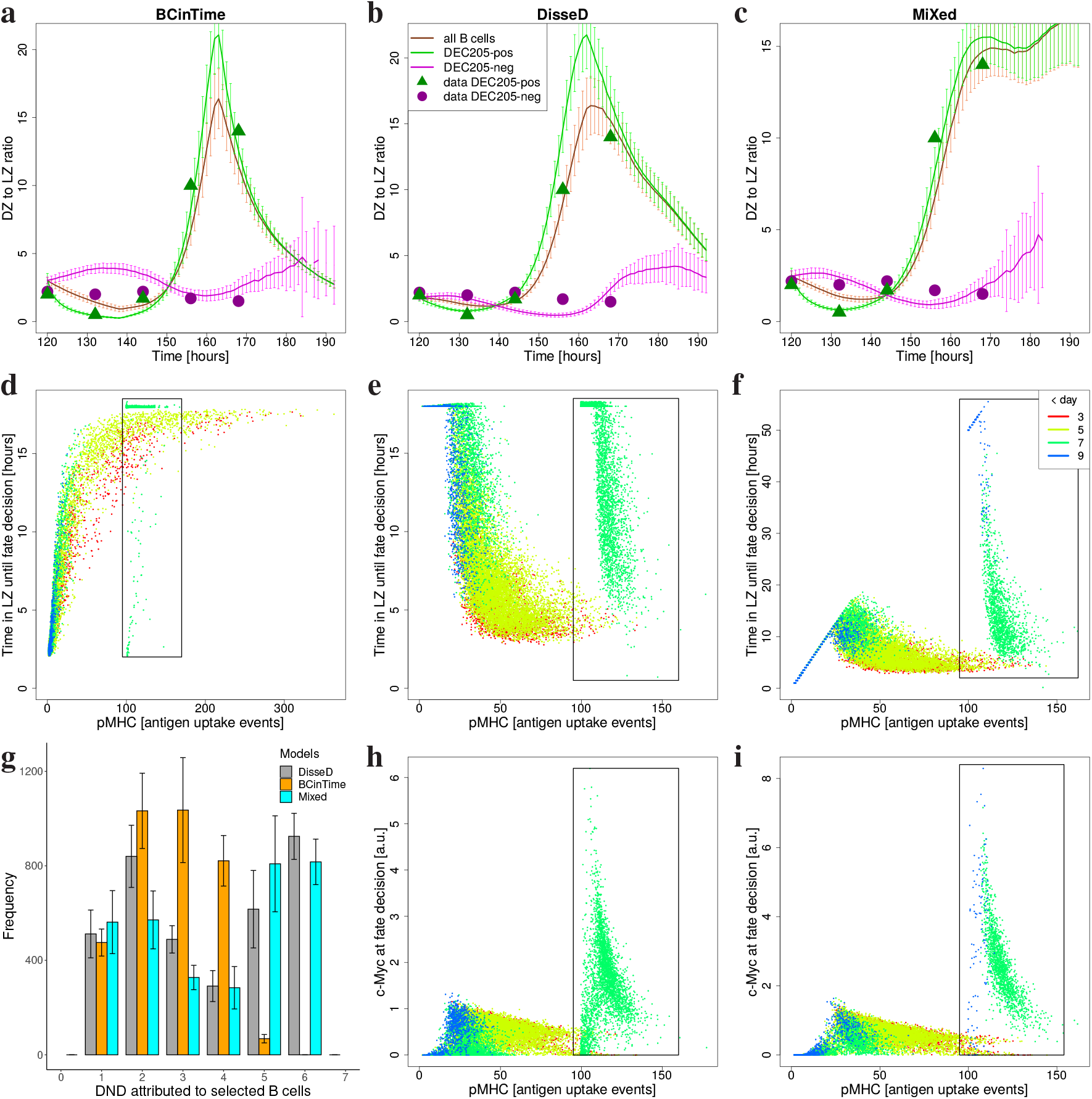
Providing antigen via the DEC205-receptor. Simulations starting from founder clones at a distance of two mutations to the optimal clone and with 20% DEC205-positive BCs (80% negative). anti-DEC205-OVA is injected *in silico* at day 5 post GC onset, which is assumed to increase pMHC-presentation to the equivalent of 100 antigen uptake events (corresponding to a five-fold higher pMHC-presentation seen in (Victora *et al.*, 2010), compared to the normal 20 in Figure 2b) in all unselected DEC205-positive BCs in the LZ in a time window of 24 hours. **(a-c)**: DZ to LZ ratio in the three models DisseD (a), BCinTime (b), MiXed (c), and comparison to experimental data (symbols) from (Victora *et al.*, 2010). Mean and SD of 100 simulations for each theory. **(d-f,h,i)**: LZ passage time (d-f) and c-Myc (h,i) in dependence on pMHC in BCinTime (d), DisseD (e,h), and MiXed (f,i). Each dot represents one cell from a single simulation. Color codes time (see inset). Green and blue dots in rectangles are anti-DEC205-OVA targeted DEC205-positive BCs. **(g)**: Distribution of the number of BC divisions. Mean and SD of 100 simulations for each theory.

In the BCinTime-theory, LZ accumulation is a natural result of the principle that BCs get additional Tfh search time with every uptaken antigen. The amount of pMHC is increased by the *in silico* injection, and so is the LZ passage time by definition (Figure 2d, rectangle). In the case of the DisseD- and MiXed-theory, LZ accumulation can only be understood by analysis of mTOR, FoxO1, and c-Myc dynamics. Injection of anti-DEC205-OVA infers high pMHC while signals downstream of BCR are weak. Low BCR-signaling impacts on FoxO1 and mTOR (Supplementary Eqs. (4) and (11)), thus, inducing a slower up-regulation of mTOR despite high pMHC levels (Supplementary Figure 2d; Figure 3e,f, rectangles), which explains the LZ accumulation. Low BCR also delays release of the FoxO1-brake on c-Myc (Supplementary Figure 2e). As the DEC205-positive BCs continue to collect antigen from FDCs, they finally get some BCR signals. In the model, FoxO1 is more sensitive to low BCR signals than mTOR, thus, the FoxO1-brake is released first allowing for c-Myc upregulation while mTOR upregulation is kept slow (Supplementary Figure 2d,f). Because of the longer LZ passage time, the BCs have the chance to interact with many Tfh, such that BCs exhibit higher c-Myc levels at the time of selection (Figure 3h,i, rectangles, Supplementary Figure 2g,h, green lines), subsequently leading to the DZ burst (Figure 3g and Supplementary Figure 2i, grey and blue).

Consistent with the *in vivo* results (see Fig. 7G in (Victora *et al.*, 2010)), affinity maturation is stopped by anti-DEC205-OVA-injection *in silico* (Supplementary Figure 3) because BCs show high pMHC to Tfh irrespective of the actual quality of the BCR. The production of output cells in response to anti-DEC205-OVA injections was markedly enhanced in all three theories (Supplementary Figure 3g) but not to the extent seen *in vivo*, suggesting a polarization of BC fate decisions towards the plasma cell phenotype in the case of high pMHC-presentation.

### LZ passage times differ in the three theories

Migration between the DZ and LZ and back is a characteristic property of GC reactions and was quantified by photoactivation of cells in each of the GC zones (Victora *et al.*, 2010). This revealed a strong asymmetry between both directions of transmigration, which was reproduced by all three theories (Figure 4a) with a tendency to underestimate DZ to LZ migration. Note that this read-out was not fine-tuned by parameter variation and has to be considered as an independent validation of the theories.

**Figure 4:**
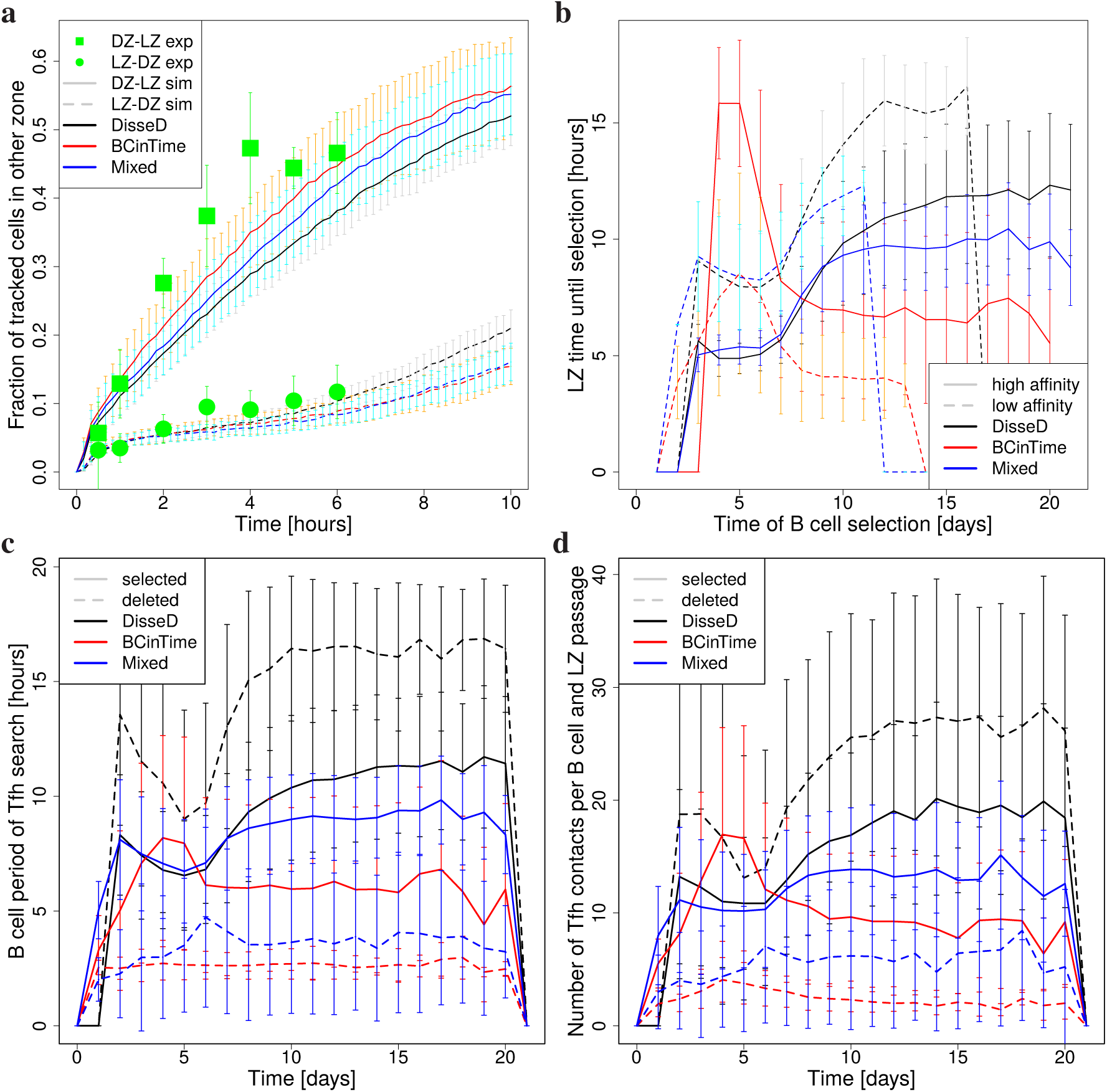
Transzone migration and LZ passage time for high and low affinity B cells. **(a):** GC simulations were mounted with high affinity founder BCs at a distance of two mutations to the optimal clone, which reflects the experimental setup based on B1-8^hi^ BCs with high specificity for NP (Victora *et al.*, 2010). BCs in the DZ (full lines) or in the LZ (dashed lines) were marked *in silico* and tracked. The fraction of marked BCs found in the opposite zone was measured over time and is shown for the three theories (colors). Mean and standard deviation of 100 simulations. Experimental data reproduced from (Victora *et al.*, 2010). **(b):** The LZ passage time (with random founder BCs) is distinguished for high (full lines) and low (dashed lines) affinity BCs in the three theories. High affinity is defined as a binding probability of more that 0.6 (Supplementary Eq. (1)). **(c):** Same as (b) for selected and deleted BCs. **(d):** Number of interactions with Tfh per BC in one LZ passage period. In B-D, mean and standard deviation over all BCs at day 7 of a representative GC simulation for each theory.

The LZ passage time, i.e. the time a BC takes from differentiation to the LZ phenotype until initiation of back differentiation to the DZ phenotype, behaves different in the different theories. Intuitively, one may think that high affinity BCs are more efficient in receiving selection signals and, thus, would have a shorter LZ passage time. This is, indeed, what is observed in the DisseD-theory (Figure 4b, black lines), where in the phase of strong selection low affinity take 8 hours while high affinity BCs manage to get through the LZ state in 5 hours. Later, when antigen becomes limiting, both times get longer. In contrast, in the BCinTime-theory (red lines), in which antigen uptake prolongs the LZ passage time, this relation is inverted and high affinity BCs take longer to pass the LZ. When antigen becomes limiting, the LZ passage times get shorter. In the MiXed-theory (blue lines), the high and low affinity BCs behave similar to the DisseD-theory because high affinity cells reach the mTOR-threshold earlier. The rise in LZ passage times when antigen gets sparse is less pronounced. This relation of LZ passage time to affinity is also reflected in the amount of collected pMHC (Figure 3d-f, ignoring green and blue dots in the rectangles). A quantitative evaluation of the correlation of the LZ passage time with affinity of all BCs at day 5 post GC onset results in Pearson correlation coefficients of −0.76 (95% confidence interval [−0.79, −0.73]) for the DisseD-, −0.73 (95% confidence interval [−0.77, −0.69]) for the MiXed-, and +0.78 (95% confidence interval [+0.76, +0.81]) for the BCinTime-theory. The correlation becomes weaker over the time of the reaction but the difference in sign is kept throughout (not shown).

Next, the LZ passage time of deleted and selected BCs is compared (Figure 4c). While deleted BCs exhibit a rather short LZ passage time in the two theories with antigen-uptake-dependent LZ time, i.e. in the BCinTime- and MiXed-theory, the LZ passage time is long in the DisseD-theory. Because of weak BCR signaling, mTOR is not upregulated quickly in the BCs (see Supplementary Eq. (11)) such that the fate decision threshold is reached at rather late times or not reached at all if the BCs hit an imposed maximum LZ passage time. Thus, the LZ passage time of deleted BCs is diametrally different in the DisseD-theory. Consequently, the number of interactions with Tfh per deleted BC, which is in the range of 10-20 per LZ passage for selected BCs, is reduced to a few contacts in the BCinTime- and the MiXed-theory, while it is increased to up to 25 but inefficient contacts in the DisseD-theory (Figure 4d).

### BCinTime-theory is more sensitive to antigen titration

*In vivo* GC reactions turned out rather robust against variations of the amount of antigen used in immunization (Vora *et al.*, 1997; Hannum *et al.*, 2000). Reducing antigen by a factor of five *in silico* reduced the peak GC population to one third in all theories (Supplementary Figure 4a). Increasing antigen by a factor of five retarded GC shut-down (Supplementary Figure 4b). This also induced a 2-3-fold peak population in the BCinTime-theory, but only a small increase in the DisseD- and the MiXed-theory (Supplementary Figure 4b). Thus, the GC reaction in the BCinTime-theory is more sensitive to the total amount of antigen and less sensitive to Tfh (see also Figure 2b,e). Despite substantial changes in the total population kinetics, DND remains similar in all high and low antigen settings in all theories (Supplementary Figure 4c,d) and is a robust GC property (Gitlin *et al.*, 2014).

### All theories are consistent with mTOR overexpression experiments

mTOR over-expressing BCs when co-existing with normal BCs in GCs have a competitive disadvantage, exhibit a larger DZ to LZ ratio, and hardly affinity mature (Ersching *et al.*, 2017). This was observed by mixing BCs that express Tsc1 normally with BCs deficient in Tsc1. Tsc1 is an inhibitor of mTOR and the Tsc1-deficient BCs express more mTOR. This experiment was replicated in the DisseD- and the MiXed-theory, which both explicitly model mTOR dynamics, by imposing a larger starting level of mTOR in BCs at LZ entry. The simulations revealed the same qualitative behavior (Supplementary Figure 4e,f). As *in vivo*, the DZ to LZ ratio was enhanced to 3-5 in the Tsc1-deficient BCs around day 4-5 post GC onset, which corresponds to day 7 post immunization. These BCs disappeared faster from the GC than *in vivo*. Affinity maturation was inhibited in Tsc1-deficient BCs similar to the *in vivo* case (data not shown). In the BCinTime-theory, mTOR is replaced by an unspecified differentiation signal. Imposing a larger starting value of this signal is a phenomenological representation of Tsc1-deficiency and induces the same phenotype as described above (Supplementary Figure 4e,f; red lines). All together, release of mTOR inhibition *in silico* induced a GC phenotype consistent with *in vivo* observations. This supports the credibility of the proposed signaling models.

### Clonal dominance gradually differs in the theories

It was shown before that GCs exhibit a great diversity of how strong founder BCs dominate a reaction (Tas *et al.*, 2016): Either a single GC founder clone bursts and then dominates the GC reaction or many founder clones co-exist during the whole GC reaction. This was revealed by staining GC BCs with one of ten random colors (confetti mice), which is kept in all progeny, and subsequently observing the dominance of the color in the course of the reaction. The same staining procedure was applied *in silico*. As in experiment, a large diversity of color dominance is found (Figure 5a-c) in all three theories. The diversity of color dominance is rising faster in the DisseD- and the MiXed-theory and the increase in the baseline dominance observed *in vivo* is only recapitulated in the BCinTime- and the MiXed-theory.

**Figure 5:**
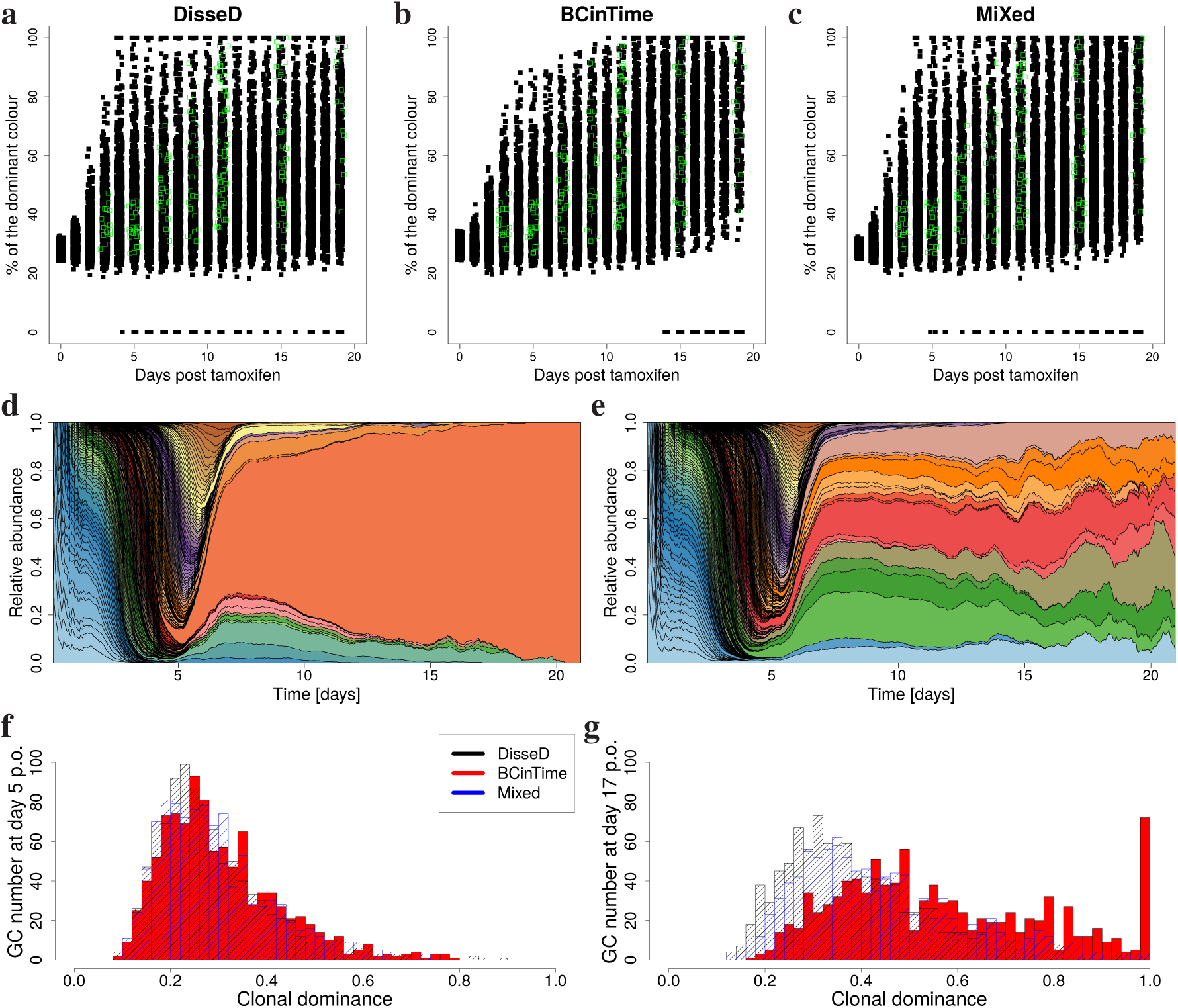
Clonal dominance. **(a-c):** Experiment with confetti mice, in which one of 10 colors is activated by tamoxifen. Data (green open squares) reproduced from (Victora *et al.*, 2010). Following the experimental protocol, one of ten colors is attributed to GC BCs with given probabilities (Table 1 of (Meyer-Hermann *et al.*, 2018)). Colors are inherited by daughter cells, the dominance of colors is monitored and reported at different GC time points for the DisseD-(a), the BCinTime-(b), and the MiXed-theory (c). **(d-e):** Muller diagram showing the relative abundance of founder clones (colors) over the course of the GC reaction (time resolution 1 hour). Two extreme cases are shown for the MiXed-theory. See Supplementary Figure 5a-d for the other theories. **(f-g):** Each founder clone and its progeny is tracked *in silico*. The largest fraction of BCs stemming from one clone is monitored. The number of GCs among 1000 GCs with a most dominant clone of particular dominance (horizontal axes) is depicted at days 5, 17 post GC onset (p.o.). More time points in Supplementary Figure 5.

While it was shown before that color and clonal dominance correlate (Meyer-Hermann *et al.*, 2018), simulations offer the possibility of following the abundance of every clone, which corresponds to sequencing every GC cell at multiple time points of the GC reaction. Both, clonal bursts leading to monoclonal GCs and long-term co-existence of many clones in a single GC were observed *in silico* (Figure 5d,e), irrespective of the theory (Supplementary Figure 5a-d).

The clonal dominance distribution over many GCs unraveled gradual differences between the theories (Figure 5f,g; Supplementary Figure 5e-j): Clonal dominance starts low and increases to levels around 30% dominance by day 5 in all theories. Later on, only a few monoclonal GCs evolve in the DisseD-theory and the dominance level is kept in the same range. In the MiXed-theory, the dominance level further increases to all possible values and a small subset of GCs exhibit clonal bursts associated with monoclonal GCs. In contrast, in the BCinTime-theory monoclonal GCs after clonal bursts appear in 20% of the GCs. The other 80% GCs are distributed over a wide range of dominance but with an overall increased dominance level. This implies that the three theories can be distinguished in experiment by the frequency of monoclonal GCs. Note that the absolute frequencies of monoclonal GCs depend on the quality of the GC founder BCs (Supplementary Figure 6).

## Discussion

### Separation of selection and activation signals

Three theories of GC BC selection were proposed on the level of intracellular antigen processing and molecular signaling. Previous GC selection models did not allow to capture all aforementioned experimental results which is now achieved by all three theories presented here (see the summary in Table 1). In particular, the physiologically extreme anti-DEC205-OVA experiments (Victora *et al.*, 2010) now appear as a natural result of the selection mechanisms at work. The contradiction between the assumed selection threshold and the DND of the recycled BCs was resolved in all three theories by the separation of selection and activation signals. In the BCinTime-theory, the time of selection is determined by the amount of antigen-uptake, while in the DisseD- and MiXed-theories it is determined by a critical mTOR level. DND is determined by the integrated Tfh signals in all theories. The separation of those signals complements the known spatial separation of division and selection into DZ and LZ and solves a long-standing inherent problem of GC selection models.

In the DisseD- and the MiXed-theory, the time point of selection of BCs is determined by BCR- and Tfh-dependent mTOR upregulation. mTOR upregulation was found in experiment, as indicated by downstream phosphorylation of S6 (Luo *et al.*, 2018). The theory is consistent with the finding that mTOR is necessary for recycling of GC BCs but block of mTOR after selection does not impact on the number of divisions (Ersching *et al.*, 2017). The exact rule of how to integrate BCR and Tfh signals is not known. In the theory, mTOR can be upregulated by each of both signals, such that sufficiently strong BCR signals alone might induce mTOR. As BCR signals are assumed a pre-requisite of mTOR upregulation by Tfh signals (Supplementary Eq. (11)), BCR and Tfh signals synergize with each other.

According to the theories, the number of divisions is determined by Tfh-dependent c-Myc levels acquired in the mTOR-determined LZ passage time. This allows for a DND as observed *in vivo* (Gitlin *et al.*, 2014). c-Myc is a powerful driver of anabolic metabolism and cell growth (Wilhelm *et al.*, 2016), which was associated with selection and recycling of GC BCs (Dominguez-Sola *et al.*, 2012), but without distinction of selection versus DND. c-Myc is essential for GC maintenance (Calado *et al.*, 2012) and acts as a timer for BC division (Heinzel *et al.*, 2017; Finkin *et al.*, 2019). The assumed fast c-Myc decay increases the selection pressure because less frequent encounter of Tfhs induces loss of c-Myc in between two interactions with Tfhs. Thus, only BCs with a high frequency of productive B-Tfh-interactions, i.e. those with high pMHC, manage to achieve higher c-Myc scores. Without c-Myc decay, low affinity BCs may acquire higher c-Myc scores because of their longer LZ passage time.

In the simulation, c-Myc upregulation is inhibited by FoxO1 (see (1 − *H*_C_(*F*)) in Supplementary Eq. (10)). The FoxO1-brake on c-Myc was released in dependence on BCR signals, as suggested by data (Luo *et al.*, 2018). Indeed, activation of Akt and PI3K downstream of BCR (Tzivion *et al.*, 2011) translocate FoxO1 to the cytosol (Burgering, 2008), which leads to FoxO1 inactivation (Brunet *et al.*, 1999). This BCR-dependent mechanism further increases the competitive advantage of high affinity BCs. Release of the FoxO1-brake opens the path for Tfh-dependent upregulation of c-Myc via CD40 ligation and the NF*κ*B pathway (Luo *et al.*, 2018). Therefore, BCR and Tfh signals synergize to upregulate c-Myc.

In the BCinTime-theory, the separation of selection and division signals is realized in a phenomenological language. The LZ passage time and consequently the selection signals are now strongly coupled to antigen-uptake rather than to Tfh signals. This infers a higher sensitivity to variations of antigen concentration (see below).

### The theories differ in the GC clonal composition

The current experimental data situation does not allow to determine a clear preference for any of the three theories. The diversity of clonal dominance in different GCs is found in all theories (Figure 5a,b). This relies on a higher degree of non-linearities in the DisseD- and the MiXed-theory (see Supplementary Table 1) because of more subtle differences in the achieved c-Myc levels, which still have to induce differential DND. The degree of non-linearities was, in particular, determined by the strong division burst observed in response to the anti-DEC205-OVA experiment (Victora *et al.*, 2010). Comparison of *in vivo* and *in silico* experiments using the Brainbow-allele, which allows to track BC clones and their progeny through the GC reaction (Tas *et al.*, 2016), favors the BCinTime-theory (Figure 5c-e). However, the frequency of mono-colored GCs depends on the choice of the set of founder BCs. Further, the number of GCs investigated *in vivo* is not sufficient to draw a conclusion. Sequencing of all GC BCs at late time points of the reaction would allow to better distinguish the theories (Figure 5f,g).

### High- and low-affinity B cells differ in their LZ passage times

A clear cut distinction of the theories is possible by a measurement of the LZ passage times of high and low affinity BCs (Figure 4b). In the BCinTime-theory, high affinity BCs stay longer in the LZ than low affinity BCs. This is inverted in the DisseD- and the MiXed-theory, where the LZ passage time is determined by mTOR in dependence on both, BCR and Tfh signals. In high affinity BCs, higher BCR-activation induces faster mTOR upregulation (Supplementary Eq. (11)), productive B-Tfh contacts become more frequent because of Tfh polarization to the BCs with highest pMHC density (Depoil *et al.*, 2005), and each contact provides more intense signals to the BC (see *H*_I_(*t*)*T* (*p*) in Supplementary Eq. (11)) which reflects synapse intensity and a positive feedback loop involving ICOSL upregulation in BCs (Liu *et al.*, 2015), together, inducing faster selection. As in the MiXed-theory, BCs have to collect antigen for survival (as in the BCinTime-theory) but the ultimate LZ passage time is determined by mTOR (as in the DisseD-theory), the drastic increase of LZ passage times found in the DisseD-theory in the late reaction was not observed in the MiXed-theory. A possible set-up for a decisive experiment is the usage of two BC lines with different affinity for the antigen in a AID-KO system with suppressed mutations. Tracking of those cells through the LZ would reveal the LZ passage time of both BC types. In a measurement around day 10 post immunization, a two-photon setting lasting for 10 hours would be needed. Thereby, the relative LZ passage times of high and low affinity BCs are more informative than the absolute times, because the absolute times vary in the course of the GC reaction while the relative times are kept throughout the reaction.

### GCs are more permissive to B cells with low BCR signals in the DisseD-theory

In the DisseD-theory, the integration of BCR and CD40 signals for upregulation of c-Myc (Supplementary Eq. (11)) includes dynamics from a positive feedback loop between ICOSL and CD40 (Liu *et al.*, 2015), and a BCR-dependent release of the FoxO1-brake. In reality, BCR signals release the brakes on Tfh-mediated signals via the NF*κ*B-pathway (Luo *et al.*, 2018), which is phenomenologically included in the FoxO1-brake. A direct impact of BCR signals onto c-Myc upregulation was not considered. Still, in the DisseD-theory, BCR signals have to be strong enough (as might be monitored by Syk-phosphorylation in reality), such that Tfh signals can induce a relevant effect on c-Myc upregulation. This implies that low affinity BCs, i.e. inefficient antigen collectors, eventually get selected because of slow but persistent mTOR activity, but receive poor division signals (i.e. low DND). This is consistent with data showing low affinity survivors in GCs (Kuraoka *et al.*, 2016) or the persistence of unrelated BCs in the GC reaction (Silver *et al.*, 2018). More generally speaking, GCs in the DisseD-theory are more permissive than in the other theories.

### FoxO1 is more sensitive to BCR signals than mTOR

The assumption that c-Myc upregulation relies on BCR signals in the DisseD-theory might be considered in contradiction to the anti-DEC205-OVA experiments (Victora *et al.*, 2010), which showed a BCR-signaling independent high DND. However, the prolonged LZ phase still allows for an accumulation of c-Myc signals *in silico* despite the inhibition of Tfh-signals in BCs with low BCR signals. Also in reality, despite removal of Syk-signaling downstream of BCR, partial upregulation of c-Myc was still found (see Figure 7 in (Luo *et al.*, 2018)). In the model, anti-DEC205-OVA-targeted BCs continue interacting with FDCs, such they receive a minimum of BCR signals. c-Myc upregulation becomes possible provided FoxO1 inhibition is more sensitive to BCR signals than mTOR (as guaranteed by the condition *K*_*F*_ < *K*_*R*_; see Supplementary Methods). Then, low BCR signals prolong the LZ phase by slow mTOR upregulation but allow for c-Myc signals because of moderate release of the FoxO1-brake.

### Mechanisms controlling the LZ passage time

In the BCinTime- and the MiXed-theory, each antigen-uptake event is associated with a prolongation of the time period of collecting signals from FDC and Tfh in the LZ. Mechanistically, the determination of the LZ time has to be independent of BCR signals because providing the antigen via the DEC205-receptor also induces a prolongation of the LZ passage time *in vivo* (Victora *et al.*, 2010), as recapitulated by the BCinTime-theory (Figure 3a). The prolongation of the LZ time cannot rely on the antigen-processing apparatus either, because providing an irrelevant antigen via the DEC205-receptor does not induce a longer LZ passage time (Victora *et al.*, 2010). A possible scenario is implemented in the MiXed-theory, where the BC life time is prolonged with antigen uptake, reflecting the old idea of suppression of pre-activated apoptosis by antigen uptake (MacLennan, 1994). The inhibition of centrocyte apoptosis is believed to depend on the interaction with FDCs and Tfh (Lindhout *et al.*, 1995; Brandtzaeg, 1996; Tew *et al.*, 1997; Hollmann & Gerdes, 1999; Hur *et al.*, 2000; van Eijk *et al.*, 2001). In the present context, only the interaction with Tfh can be responsible of suppressing apoptosis because the prolongation of the LZ phase happens independent of interactions with FDCs. Alternatively, rather than targeting apoptosis, Tfh might prevent acquisition of the DZ phenotype in highly pMHC-loaded BCs by suppression of FoxO1 via activation of Akt (Luo *et al.*, 2018). Activation of Akt would be stronger in high-pMHC BCs, thus, enforcing higher levels of mTOR to suppress Akt (Ersching *et al.*, 2017) and allow for FoxO1 up-regulation, thus, delaying recycling and acquisition of the DZ-phenotype.

### Speculation on adaptive control by Tfhs

Antigen titration revealed a robustness of the number of divisions per selection round despite rather different population kinetics. The BCinTime-theory was most sensitive to the amount of antigen presented on FDCs, while the MiXed- and the BCinTime-theory more rely on Tfh help. In view of the robustness of GC reactions kinetics to changes in the levels of immune-complex deposition on FDCs (Vora *et al.*, 1997; Hannum *et al.*, 2000) one may speculate whether Tfh help is adaptive to the overall level of pMHC-levels detected on BCs. In such a model, Tfh signaling intensity would be gauged by previously seen pMHC-densities, thus, allowing for proper support of BCs in a framework of low antigen availability.

### Evolutionary reasons for signal separation

With computer simulations the separation of signals for the frequency of selection in the LZ and for the strength of divisions in the DZ was identified as a concept of information processing in GCs that reconciles seemingly contradicting experimental data. Separation of signals is a well-known concept, e.g. in digital audio signals, which enables independent modulation of different parts of information. Evolutionary, this allows for a larger diversity of BCs surviving GC selection but limited by their low division potential, thus, keeping the focus on the need for high affinity antibodies in the acute situation. The survival of low affinity cells is reflected in the diversity of clonal dominance in GCs and is critical for flexibility of the immune system facing fast mutating pathogens.

### Evolutionary differences of the three theories

From the evolutionary perspective, the DisseD- and the MiXed-theory appear advantageous. High affinity cells are selected faster, which might be important in view of a time-critical fight against an infection. Robustness against variations of antigen amount is higher, which makes the immune system less vulnerable to immune evasion strategies of slowly expanding pathogens. Less GCs are monoclonal inferring a better preparation for future infections with mutated pathogens by a diverse memory repertoire. The presented computer simulations predict testable read-outs that allow distinguishing the three proposed theories and by that to shed light on mechanisms at work in real GCs and paving the ground of improved vaccination protocols. I hope to stimulate further experimentation along these lines.

## Methods

The details of the presented computer simulations are described in the Supplementary Material. The software was implemented in C++11 on a Linux platform. Figures were generated from the simulation read-out using R.

## Acknowledgement

MMH thanks Mark J Shlomchik, and Gabriel D Victora for fruitful discussions, Sahamoddin Khailaie for preparation of Figure 1 and revising the manuscript, and Sebastian C Binder, Coraline Mlynarczyk, and Ari M Melnick for revising the manuscript. MMH was in parts supported by the Human Frontier Science Program (RGP0033/2015).

## Author contribution

MMH developed the research idea and the approach, programmed the code, interpreted the results and wrote the manuscript.

## Supplementary material

**Table 1:** Theory-specific parameter values. Equation numbers and symbols refer to the Supplementary Methods.

| model name | BCinTime | DisseD | MiXed |
| --- | --- | --- | --- |
| antigen per FDC | 3000 | 10000 | 10000 |
| minimum Ag-uptake | 1 | 2 | 2 |
| CC-CB differentiation delay [h] | 7 | 6 | 6 |
| Tfh number | 200 | 100 | 100 |
| LZ passage time given by | Eq. (8) | Eqs. (5) and (11) | Eqs. (5) and (11) |
| $K_R$ | 4 | 20 | 28 |
| $n_R$ | 2.4 | 2 | 2 |
| Tfh-B-interaction time | Gauss | $p$ | $p$ |
| min-value [h] | - | 0 | 0 |
| max-value [h] | - | 0.3 | 0.2 |
| K-value [$p$] | - | 20 | 15 |
| Hill-coefficient | - | 2 | 2 |
| $T(p)$ | 1 | $T(p)$ | $T(p)$ |
| min-value (factor) | - | 0 | 0 |
| max-value (factor) | - | 3 | 4 |
| K-value, pMHC(factor=1) | - | 15 | 15 |
| Hill-coefficient | - | 2 | 2 |
| $C$ | Eq. (9) | Eq. (10) | Eq. (10) |
| $C_{\text{threshold}}$ | 0.3 | 0.4 | 0.2 |
| $\ln(2)/\gamma_C$ [h] | - | 3 | 3 |
| $n_{\text{DND}}$ | 1.3 | 4.0 | 3.3 |
| $C$ for DND = 2 | 0.8 | 0.7 | 0.82 |

**Supplementary Figure 1:**
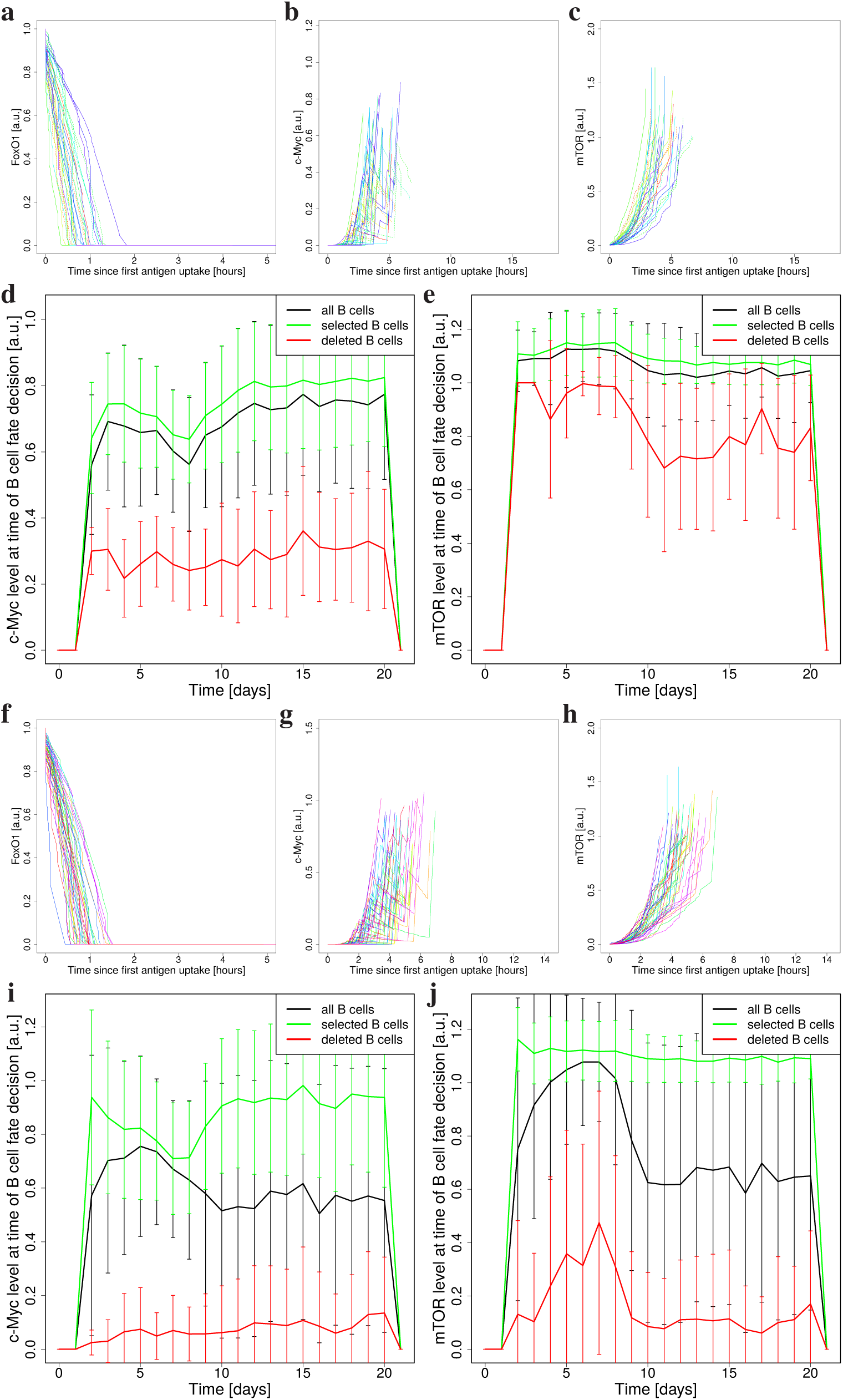
Dynamics of key molecules. **(a-c,f-h)**: The time course of FoxO1 (a,f), c-Myc (b,g), and mTOR (c,h) for 20 randomly selected BCs at day 7 post GC onset(a-c, DisseD; f-h, MiXed). **(d,e,i,j)**: Mean and standard deviation of c-Myc (d,i) (selection threshold 0.4 for DisseD; 0.2 for MiXed) and mTOR (e,j) (threshold for fate decision is 1) of all LZ-BCs over the course of the GC reaction (d,e, DisseD; f-h, MiXed).

**Supplementary Figure 2:**
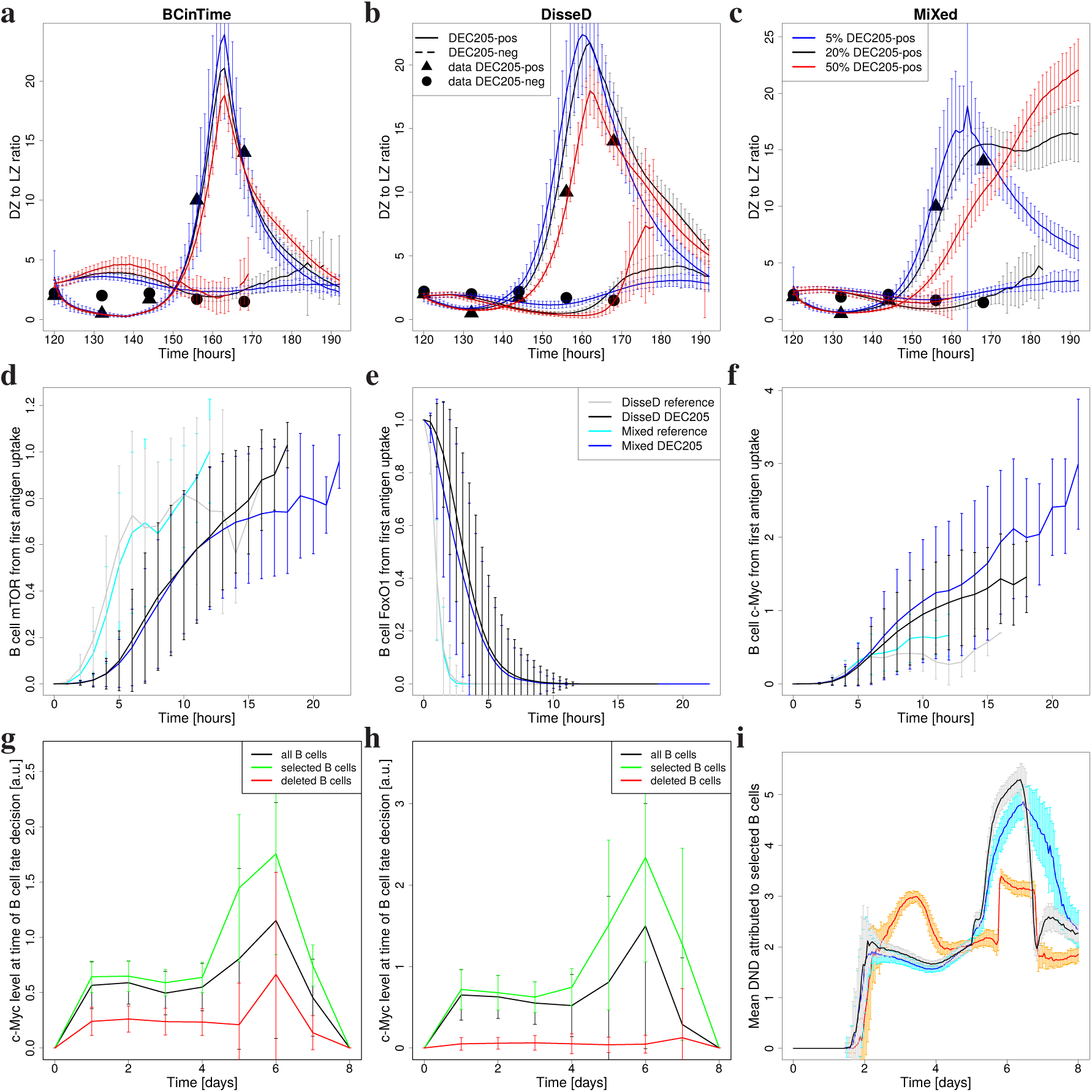
GC dynamics upon anti-DEC205-OVA injection as in Figure 3a-c. **(a-c)**: DZ to LZ ratio with 5% (blue), 20% (black), and 50% (red) DEC205-positive BCs. **(d-f)**: Dynamics of mTOR, FoxO1, and c-Myc in DisseD (grey and black) and MiXed (cyan and blue) with (black and blue) and without (grey and cyan) anti-DEC205-OVA injection. Time zero is the time of first antigen acquisition of each BC. Mean and SD of all selected BCs at day 6. **(g,h)**: c-Myc over time in DisseD (g) and MiXed (h). Mean and SD of all LZ BCs in a single simulation. **(i)**: DND attributed to selected BCs. 20% DEC205-positive BCs in (d-i). Mean and SD of 100 simulations in (a-c,i).

**Supplementary Figure 3:**
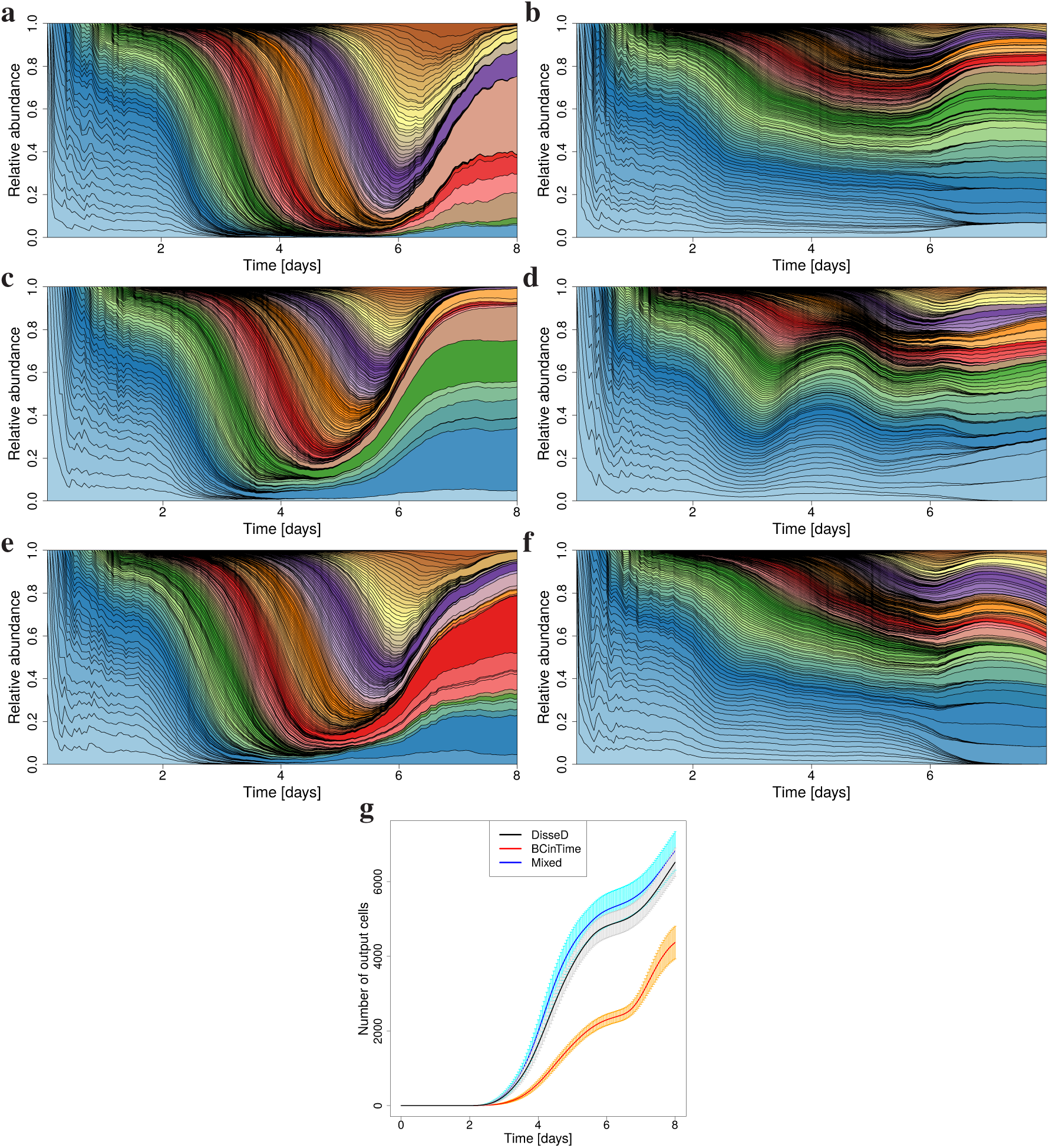
Anti-DEC205-OVA injection stops affinity maturation and accelerates output. **(a-f)**: Muller diagrams showing the evolution (time resolution 1 hour) of the relative clonal abundance in single GC simulations without (a,c,e: random founder clones) or with anti-DEC205-OVA injection *in silico* at day 5 (b,d,f). Founder clones at a distance of two mutations to the optimal clone. 20% DEC205-positive BCs. Results for the DisseD-(a,b), BCinTime-(c,d), and MiXed-theory (e,f). The progressively increasing clonal dominance is suppressed by the injection. **(g)**: Total amount of generated output. Mean and SD of 100 simulations.

**Supplementary Figure 4:**
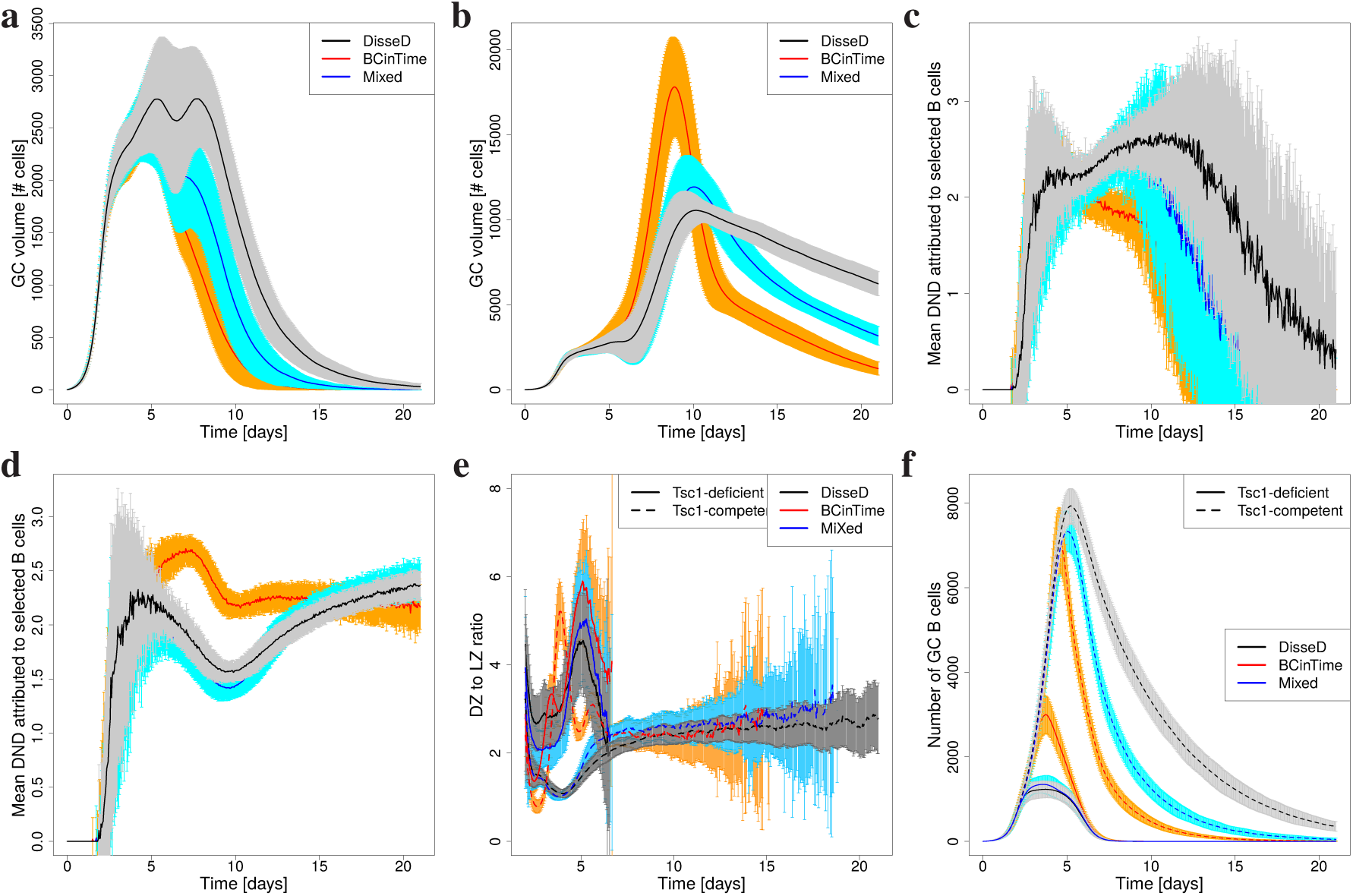
Antigen titration and mTOR over-expression experiments. **(a-d)** The total amount of antigen distributed on FDC sites is reduced (a,c) or increased (b,d) five-fold with respect to Figure 2. **(e-f)** At the time of differentiation of DZ-BCs to the LZ phenotype mTOR is not set to zero but to 80% (in the BCinTime-theory the unspecified differentiation signal is set to 50%) of the value needed to induce fate decision. The GC simulation is started with 50% of the founder cells sensitive to this modulation. As in [Ersching *et al.*, 2017], GC founder cells are assumed high affinity, i.e. at a distance of two mutations from the optimal clone. Mean and standard deviation of from 100 simulations.

**Supplementary Figure 5:**
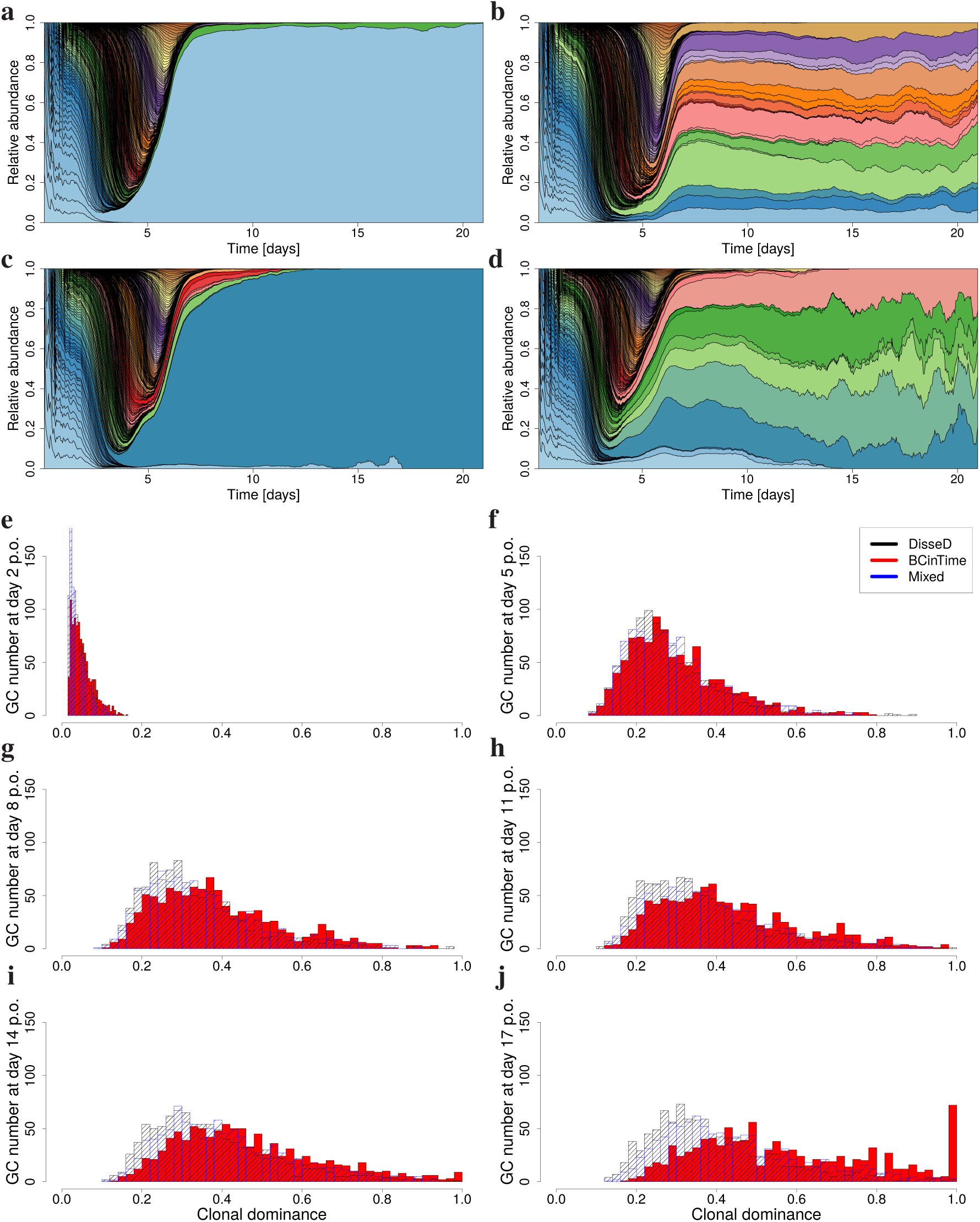
Clonal dominance with free founder cells. Extension of Figure 5: **(a-d)**: Muller diagram showing the relative abundance of founder clones (colors) over the course of the GC reaction (time resolution 1 hour). Two extreme cases are shown for the DisseD-(a,b) and the BCinTime-theory (c,d). See Figure 5 for the MiXed-theory. **(e-j)**: Each founder clone in 1000 GC simulations is tracked and the degree of dominance in terms of how many GC-BCs stem from a particular founder clone is monitored. The number of GCs among the 1000 GCs with a most dominant clone of particular dominance (horizontal axes) is depicted at days 2, 5, 8, 11, 14, 17 post GC onset (p.o.).

**Supplementary Figure 6:**
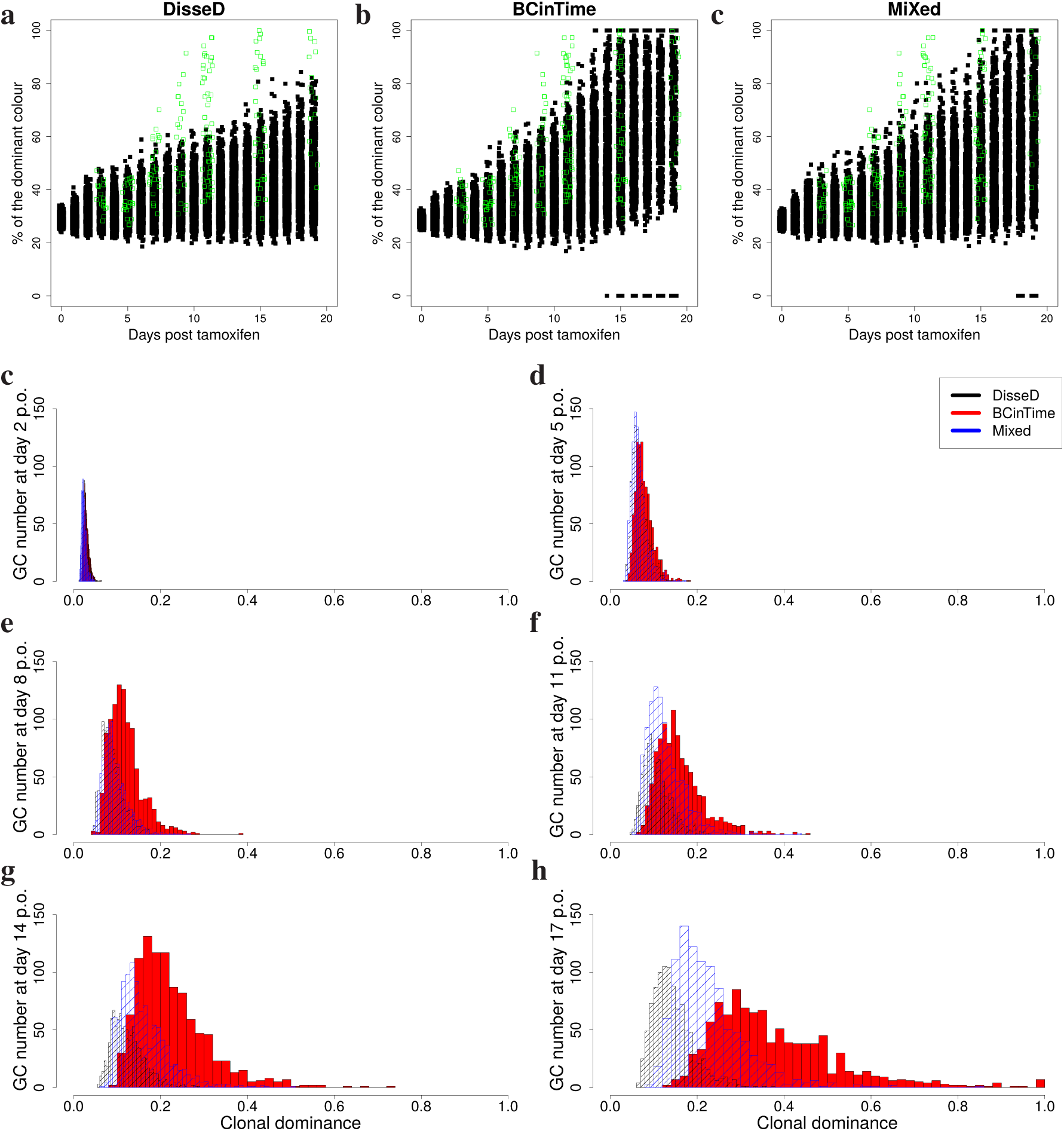
Clonal dominance with restricted founder cells. Same analysis as in Figure 5 but instead of random choice of founder cells, the founder cells are restricted to have a mutation distance to the optimal clone of 5 or 6 mutations. **(a-c)**: Experiment with confetti mice, in which one of 10 colors is activated by injection of tamoxifen. Data reproduced from [Victora *et al.*, 2010] (green open squares). Following the experimental protocol, one of ten colors is attributed to GC-BCs with probabilities as described in Table 1 of [Meyer-Hermann *et al.*, 2018]. Colors are inherited by daughter cells, the dominance of colors is monitored and reported here for different time points of the reaction for the DisseD-(a), the BCinTime-(b), and the MiXed-theory (c). **(d-i)**: Each founder clone in 1000 GC simulations is tracked and the degree of dominance in terms of how many GC-BCs stem from a particular founder clone is monitored. The number of GCs among the 1000 GCs with a most dominant clone of particular dominance (horizontal axes) is depicted at days 2, 5, 8, 11, 14, 17 post GC onset (p.o.), panels in reading order.

**Supplementary Figure 7:**
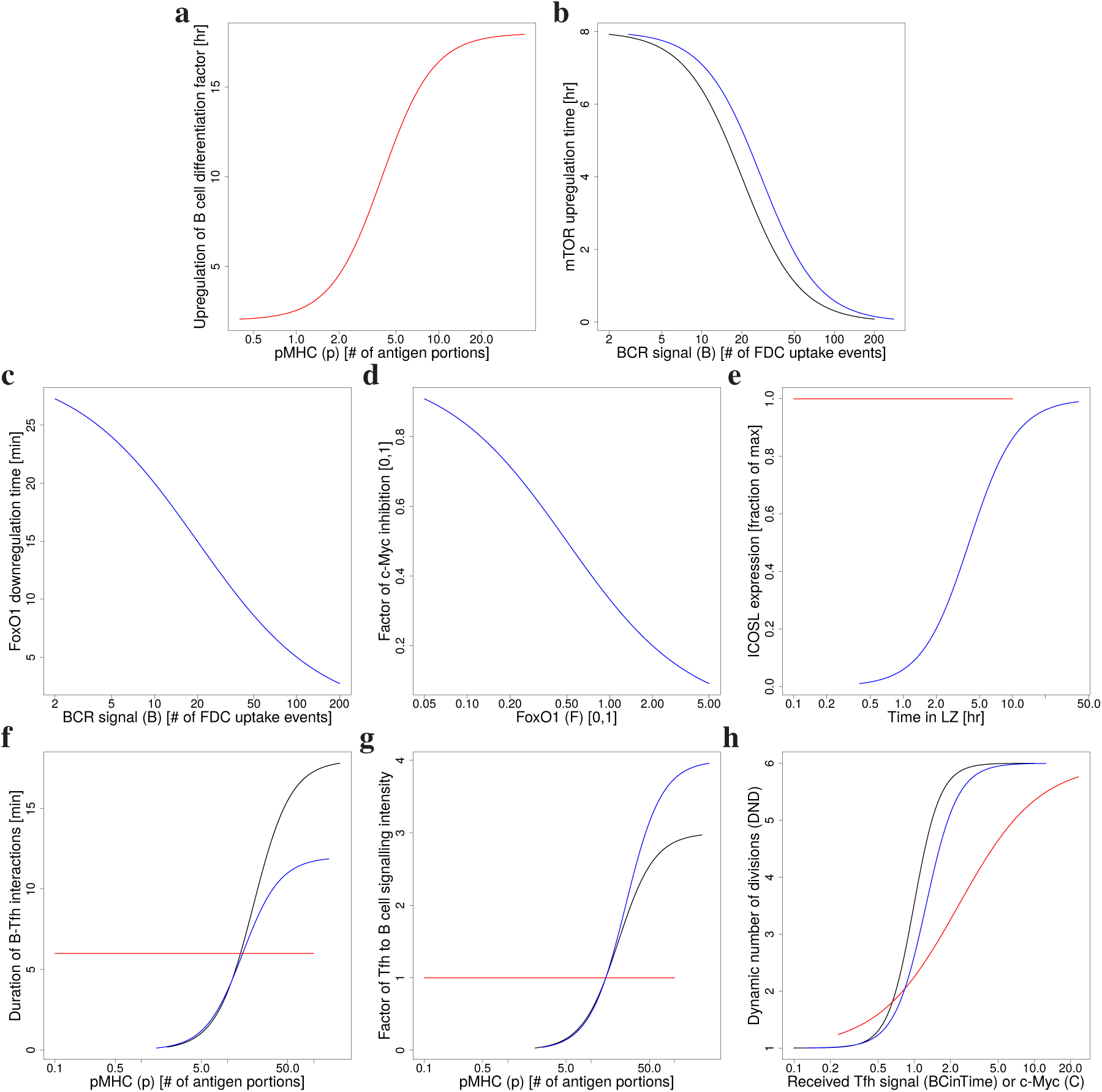
Hill functions of B cell selection. The Hill-function used in the DisseD-(black), BCinTime-(red), and MiXed-theory (blue) as defined in the Methods section. If only a blue line is shown, the black line is identical. **(a)** Inverse upregulation rate of signal *R* in Eq. (8) in dependence on presented pMHC *p* (only in the BCinTime-theory); **(b)** inverse upregulation rate *α*_R_*H*_R_(*B*) of mTOR *R* in Eq. (11) in dependence on BCR-signals *B*; **(c)** inverse downregulation rate *α*_F_*H*_F_(*B*) of FoxO1 *F* in Eq. (4) in dependence on BCR-signals *B*; **(d)** factor (1 − *H*_C_(*F*)) of inhibition of c-Myc upregulation in Eq. (10) by FoxO1 *F*; **(e)** factor *H*_I_(*t*) of intensifying Tfh signals to induce c-Myc and mTOR upregulation in Eqs. (10) and (11) by ICOSL *I*; **(f)** duration of individual Tfh-BC interactions in dependence on the presented pMHC *p*; **(g)** signaling intensity factor *T* (*p*) during Tfh-BC interactions in Eqs. (10) and (11) in dependence on presented pMHC *p*; **(h)** the DND attributed to selected BCs in Eq. (12) in dependence on *C*, i.e. received Tfh signals in the BCinTime-theory or c-Myc otherwise. (b-d) only defined in the DisseD- and the MiXed-theory.

## Supplementary methods

### Germinal Center simulation model

The germinal center (GC) model assumptions underlying the GC simulations are explained here and include the used parameter values. It follows [Meyer-Hermann *et al.*, 2012] with additional features [Meyer-Hermann, 2014, Binder & Meyer-Hermann, 2016, Meyer-Hermann, 2019], and adding the specificities of the DisseD-, the BCinTime-, and the MiXed-theory introduced here. Parameters differing in the three theories are listed in Table 1. Used acronyms are: DZ for dark zone, LZ for light zone, Tfh for T follicular helper cell, FDC for follicular dendritic cell, BC for B cell, BCR for B cell receptor.

#### Space representation

All reactions take place on a three-dimensional discretized space with a rectangular lattice with lattice constant of ∆*x* = 5*µm*. Every lattice node can be occupied by a single cell only.

#### Shape space for antibodies

Antibodies are represented on a four (*d* = 4) dimensional shape space [Perelson & Oster, 1979]. The shape space is restricted to a size of 10 positions per dimension, thus, only considering antibodies with a minimum affinity to the antigen. The optimal clone Φ^∗^ is positioned in the center of the shape space. A position on the shape space Φ is attributed to each BC.

The 1-Norm with respect to the optimal clone 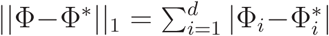, i.e. the minimum number of mutations required to reach the optimal clone, is used as a measure for the antigen binding probability. The binding probability is calculated from the Gaussian distribution with width Γ = 2.8 [Meyer-Hermann *et al.*, 2001]:

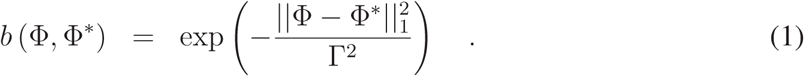

#### B cell phenotypes

Three BC phenotypes are distinguished: DZ BCs, LZ BCs, and output cells. The different phenotypes characterize the cell properties and are not meant as localization within the GC zones. DZ BCs divide, mutate and migrate. LZ BCs also migrate and undergo the different stages of the selection process. Output cells only migrate.

#### Founder cells

The model starts from a fixed number of Tfh (Table 1), 200 FDCs, 300 stromal cells, and no BC. Tfh are randomly distributed on the lattice and occupy a single node each. Stromal cells are restricted to the DZ (see section *Chemokine distribution* for their function). FDCs are restricted to the upper half of the reaction sphere, occupy one node by their soma and have 6 dendrites of 40*µm* length each. The presence of dendrites is represented as a lattice-node property and, thus, visible to BCs. The dendrites are treated as transparent for BC or Tfh migration such that they do not inhibit cell motility.

BCs enter the GC reaction with a probability per time step corresponding to a rate of 2 cells per hour (for an estimation of this value refer to [Meyer-Hermann, 2019]). New BCs are randomly positioned on free lattice nodes. The shape space position of each new BC is randomly picked from a set of 100 randomly picked shape space positions, unless stated otherwise.

#### Antigen-presentation by FDCs

Each FDC is loaded with antigen portions distributed onto the lattice-nodes occupied by FDC-soma or FDC-dendrite (Table 1). One antigen portion corresponds to the number of antigen molecules taken up by a BC upon successful contact with an FDC.

#### DZ B cell division

The average cell cycle duration of 7.5 hours of DZ BCs is varied for each BC according to a Gaussian distribution. This is needed to get desynchronization of BC division. The cell cycle is decomposed into four phases (G1, S, G2, M) in order to localize mitotic events if this is needed. Each founder BC divides a number of times before differentiating to the LZ phenotype for the first time. Six divisions was the number of divisions found in response to the extreme stimulus with anti-DEC205-OVA [Victora *et al.*, 2010, Meyer-Hermann *et al.*, 2012]. Each selected BC divides a number of times determined by the interaction with Tfh (see below, LZ B cell selection). The parameters of the interaction with Tfh are tuned such that the mean number of divisions is in the range of two [Gitlin *et al.*, 2014]. This value is required in order to maintain a DZ to LZ ratio in the range of two [Victora *et al.*, 2010, Meyer-Hermann *et al.*, 2012].

A division requires free space on one of the Moore neighbors of the dividing cell. Otherwise the division is postponed until a free Moore neighbor is available.

At every division the encoded antibody mutates with probability 0.5 [Berek & Milstein, 1987, Nossal, 1992]. This corresponds to a shift in the shape space to a von Neumann neighbor in a random direction. Upon selection by Tfh the mutation probability is individually reduced from *m*_max_ = 0.5 down to *m*_min_ = 0 in an affinity-dependent way following

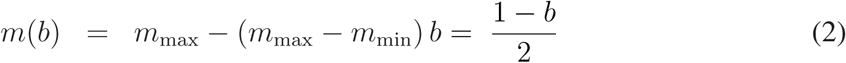

with *b* from Eq. (1) [Toellner *et al.*, 2002]. Thus, after recycling DZ BCs can acquire reduced mutation probabilities. This mechanism is motivated by the observation that BCR internalization enhances the activation of the kinase Akt [Chaturvedi *et al.*, 2011] which, in turn, suppresses activation induced cytosine deaminase (AID) [Omori *et al.*, 2006]. AID is required for somatic hypermutation, such that this provides an affinity-dependent down-regulation of the mutation frequency [Dustin & Meyer-Hermann, 2012].

BC division of BCs that previously acquired antigen and have been selected by Tfh distribute the retained antigen asymmetrically to the daughters [Thaunat *et al.*, 2012]. The model assumes asymmetric division in 72% of the cases, which is supported by experimental observations (see [Thaunat *et al.*, 2012] and Supplementary Figure S1 in [Meyer-Hermann *et al.*, 2012]). If division is asymmetric, one daughter gets all the retained antigen while the other gets none, which approximates the value of 88% found in [Thaunat *et al.*, 2012]. Mutation is suppressed in cells retaining antigen.

After the required number of divisions the BC differentiates with a rate of in 1*/*6 minutes to the LZ phenotype. All BCs that kept the antigen up to this time, differentiate to output cells, up-regulate CXCR4, and leave the GC in direction of the T zone.

#### LZ B cell selection

At the time *t* = 0 of differentiation from the DZ to the LZ phenotype, BCs are in state *unselected* and BC-specific factors are initialized to

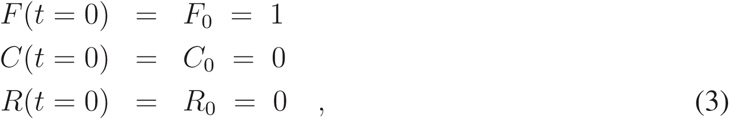

where *F*_0_ does not apply to the BCinTime-theory.

These factors are associated with FoxO1 (*F*), c-Myc (*C*), and mTOR (*R*) in the DisseD- and the MiXed-theory. In BCs not in contact to Tfh, FoxO1, mTOR and c-Myc evolve according to

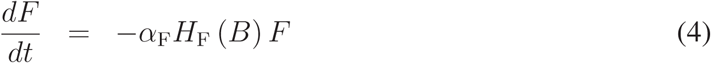

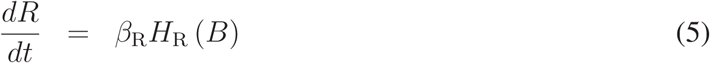

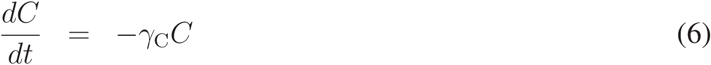

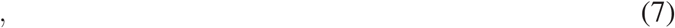

with *β*_R_ = 1/(8 hours) the mTOR production rate modulated by the BCR-dependent Hillfunction *H*_R_(*B*) (see *K*_R_ and *n*_R_ in Table 1 and Figure 7B). *α*_F_ = 2/hours is the FoxO1 reduction rate modulated by the BCR-dependent Hill-function *H*_F_(*B*) between one and zero with K-value of 20 and Hill-coefficient 1 (Figure 7C). *γ*_C_ = ln(2)/(3 hours) is the c-Myc degradation rate. *R* and *C* have different dynamics in state *Tfh-contact*.

In the BCinTime-theory, the unspecified differentiation signal (*R*) follows

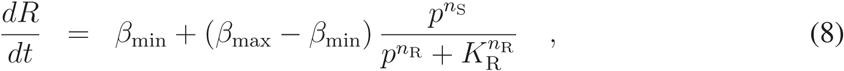

with 1/*β*_min_ = 2 hours and 1/*β*_max_ = 18 hours. Thus, the growth rate of *R* is determined by the total amount of pMHC *p* collected by the BC at any given time with *K*_R_ and *n*_R_ the pMHC amount for half max reduction of the growth rate and the associated Hill-coefficient, respectively, (see Table 1 and Figure 7A). The integrated level of Tfh signals a BC received (*C*) is defined in the state *Tfh-contact*, and *F* is not used.

LZ BCs can be in the states *unselected, FDC-contact, FDC-selected, Tfh-contact, selected, apoptotic*.

##### Unselected

LZ BCs migrate and search for contact with FDCs loaded with antigen in order to collect antigen for 0.7 hours. If an FDC soma or dendrite is present at the position of the BC, the BC attempts to establish contact to the epitope with highest affinity to the BCR (default setting). Alternatively, the BC may attempt to establish contact to the epitope of highest availability at this site. In both settings, binding is affinity dependent and happens with the probability *b* in Eq. (1). If the available number of antigen portions at the specific FDC site drops below 20 the binding probability *b* is linearly reduced with the number of available portions. If successful, the BC switches to the state *FDC-contact*; otherwise the BC continues to migrate. Further binding-attempts are prohibited for 1.2 minutes. At the end of the antigen collection period, BCs switch to the state *FDC-selected*. If a LZ BC fails to collect any antigen until this time it switches to the state *apoptotic*.

##### FDC-contact

LZ BCs remain immobile (bound) for 3 minutes [Schwickert *et al.*, 2007] and then return to the state *unselected* or *FDC-selected*, depending on from which state contact to FDC was established. The counter for the number of successful antigen uptake events *p* and the degree of BCR-signaling *B* are increased by one unit and the FDC reduces its locally available antigen portions by one. In the DisseD- and the MiXed-theory, factors depend differently on the amount of presented pMHC *p* and the degree of BCR-signaling *B*. In anti-DEC205-OVA experiments, antigen is taken up and presented without induction of BCR signals, thus, *p* in increased by 100 antigen uptake events but *B* is left unchanged. The amount of 100 is motivated by the finding that pMHC presentation is increased five-fold upon anti-DEC205-OVA injection [Victora *et al.*, 2010] together with a mean number of 20 − 30 uptake events seen under normal conditions in the simulations.

##### FDC-selected

BCs search for contact with Tfh or FDC. If they meet a Tfh they switch to the state *Tfh-contact*. If they meet a FDC they switch to the state *FDC-contact* with probability *b* in Eq. (1). If Tfhs and FDCs are neighbors of a BC, the BC binds the Tfh.

##### Tfh-contact

LZ BCs remain immobile for a time randomly sampled from a Gaussian distribution with mean 6 minutes and width 1.2 minutes (BCinTime) or for a time determined by a Hill-function in dependence on the amount of presented pMHC *p* (DisseD and MiXed), see Table 1 *Tfh-B-interaction time* and Figure 7F. During this time, the bound Tfh, which may also be bound to other BCs, polarizes to the BC with highest number of successful antigen uptakes. Polarization is determined from scratch in every time step and inertia from repolarization or intracellular organelles are ignored. If more than one BC shares the same highest pMHC presentation *p*, polarization is chosen randomly among those BCs. Only the BC to which the Tfh is polarized receives Tfh signals *C* and accumulate those.

In the BCinTime-theory, Tfh-signals follow

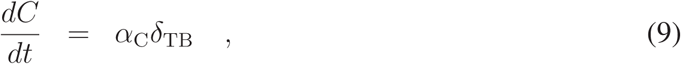

where *α*_C_ is the rate of signal acquisition during polarized Tfh-BC interactions, ensured by *δ*_TB_, which is one during polarized contact and zero otherwise). The rate is set to 1/hr without loss of generality. Thus, *C* reflects the integrated duration of Tfh-BC interactions with Tfh polarized to this BC.

In the DisseD- and MiXed-theory, Tfh signals are represented as mTOR levels as well as c-Myc levels, which are modulated by FoxO1, according to

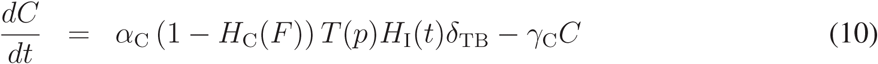

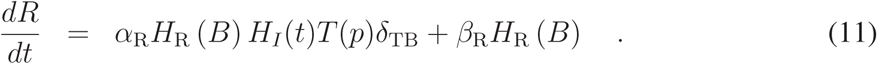

*T* (*p*) is the pMHC-dependent intensity of Tfh signaling to the interacting BC associated with CD40-signals, which is described by Hill-function (see Table 1 *T* (*p*) and Figure 7G). Signaling intensity is further modulated by the degree of ICOSL upregulation *H*_*I*_ (*t*). *H*_I_(*t*) is a Hill-function in time with values between zero and one, determined by a characteristic ICOSL-upregulation time of four hours in mice [Papa *et al.*, 2017] and Hill-coefficient two (Figure 7E).^1^ *α*_C_ and *γ*_C_ are the growth and degradation rates of c-Myc. *β*_R_ = 1/(8 hours) tunes the level of mTOR upregulation when BCs are not in contact to Tfh (see Eq. (5)), while *α*_R_ = 1/(hours) determines mTOR upregulation during productive contact with Tfh. *H*_R_(*B*) is a Hill-function describing the dependence of mTOR regulation on signals downstream of BCR (see *K*_R_ and *n*_R_ in Table 1 and Figure 7B). *H*_C_(*F*) is a Hill-function controlling the suppression of c-Myc signals by FoxO1 with values between zero and one, K-value of 0.5 and Hill-coefficient of 1 (Figure 7D).

After the binding time, the BC detaches and returns to the state *FDC-selected*. It continues to search for and bind FDC or Tfhs, where binding twice the same Tfh in a sequence is excluded. The total period of signal acquisition in the LZ is determined differently in the three theories: It finished when *R* = 1 is reached in the BCinTime-theory, when *R* = 1 or the maximum time period of 18 hours is reached in the DisseD-theory, when the LZ passage time gets longer than the number of antigen uptake events times 0.5 hours in the MiXed-theory. When one of those limits is reached during a running contact, the contact is kept active until the end of the contact time. Then, it switches to the state *apoptotic* if either *R <* 1 or if the received Tfh-signals are *C < C*_threshold_. It switches to the state *selected* otherwise.

##### Selected

LZ BCs keep the LZ phenotype for a determined time period (Table 1, CC-CB-differentiation delay) and desensitize for CXCL13, thus, perform a random walk. During that time they reenter cell cycle and progress through the cell cycle phases. Then they recycle back to the DZ phenotype with a rate of 1/6 minutes and memorize the amount of collected antigen as well as the cell cycle phase they have achieved by this time.

The number of divisions *P* (*C*) the recycled BCs will do is derived from the *C*, which reflects the amount of integrated Tfh signals (BCinTime-theory) or the c-Myc level (DisseD- and MiXed-theory), according to

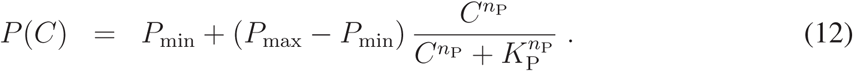

The minimum number of division is set to one (*P*_min_ = 1) in order to avoid recycling events without further division. It is limited by six divisions in the best case, which is motivated by anti-DEC205-OVA experiments in which DEC205^+/+^ BCs received abundant antigen which increased pMHC presentation to a maximum [Victora *et al.*, 2010]. The population dynamics in vivo and in silico only matched when the number of divisions was increased to six in the simulation [Meyer-Hermann *et al.*, 2012] suggesting that the strongest possible pMHC presentation to Tfh induces six divisions (*P*_max_ = 6). The Hill-coefficient *n*_P_ was theory-specific (Table 1 and Figure 7H).

The half value *K*_P_ remained to be determined, which denotes the amount of antigen collected by BCs at which the number of divisions becomes half maximal. It was fixed by the condition that for the mean level of *C* = *C*_0_ the number of divisions becomes two, as observed in [Gitlin *et al.*, 2014]:

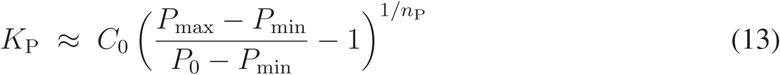

and differs for each theory (Table 1).

##### Apoptotic

LZ BCs remain on the lattice for 6 hours before they are deleted. They continue to be sensitive to CXCL13 during this time.

#### Chemokine distribution

Two chemokines CXCL12 and CXCL13 are considered. CXCL13 is produced by FDCs in the LZ with 10nMol per hour and FDC while CXCL12 is produced by stromal cells in the DZ with 400nMol per hour and stromal cell. As both cell types are assumed to be immobile, chemokine distributions were pre-calculated once and the resulting steady state distributions were used in all simulations.

#### Chemotaxis

DZ and LZ BCs regulate their sensitivity to CXCL13 and CXCL12, respectively. This is true in all BC states unless stated otherwise. All BCs move with a target speed of 7.5*µm/min*. This leads to a slightly lower observable average speed of ≈ 6*µm/min*.

BCs have a polarity vector that determines their preferential direction of migration. The polarity vector *p* is reset every 1.5 minutes into a new direction using the chemokine distribution *c* as

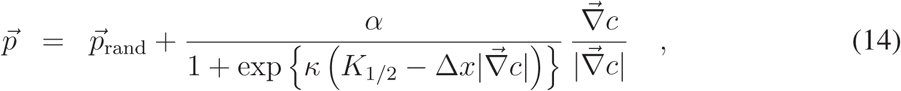

where 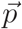_rand_ is a random polarity vector and the turning angle is sampled from the measured turning angle distribution ([Allen *et al.*, 2007] Fig. S1B). *α* = 3 determines the relative weight of the chemotaxis and random walk, *K*_1/2_ = 2 • 10^11^ Mol determines the gradient of half maximum chemotaxis weight, and *κ* = 10^10^/Mol determines the steepness of the weight increase.

BCs de- and re-sensitize for their respective chemokine depending on the local chemokine concentration: The desensitization threshold is set to 4.5nMol and 0.08nMol for CXCL12 and CXCL13, respectively, which avoids cell clustering in the center of the zones. The resensitization threshold is set at 2/3 and 3/4 of the desensitization threshold for CXCL12 and CXCL13, respectively.

BCs can only migrate if the target node is free. If occupied and the neighbor cell is to migrate in the opposite direction (negative scalar product of the polarity vectors) both cells are exchanged with a probability of 0.5. This exchange algorithm avoids lattice artifacts leading to cell clusters.

Tfh do random walk with a preferential directionality to the LZ: The polarity vector 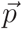 of Tfh is determined from a mixture of random walk 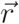 and the direction of the LZ 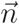 by

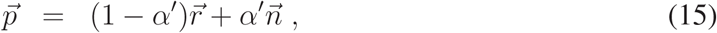

where *α*′ = 0.1 is the weight of chemotaxis. This weight leads to a dominance of random walk with a tendency to accumulate in the LZ as found in experiment. TCs migrate with an average speed of 10*µm/min* and repolarize every 1.7 minutes [Miller *et al.*, 2002].

Output cell motility is derived from plasma cell motility data to 3*µ*m per minute with a persistence time of 0.75 minutes [Allen *et al.*, 2007].

## Footnotes

1 Include the ICOSL Hill in Figure S8.

## References

Allen, C. D., Okada, T., Tang, H. L. & Cyster, J. G. (2007). Imaging of germinal center selection events during affinity maturation. Science, 315, 528–531.

Binder, S. & Meyer-Hermann, M. (2016). Implications of intravital imaging of murine germinal centers on the control of B cell selection and division. Front. Immunol. 7, 593.

Brandtzaeg, P. (1996). The B-cell development in tonsillar lymphoid follicles. Acta Otolaryngol. Suppl. (Stockh), 523, 55–59.

Brunet, A., Bonni, A., Zigmond, M., Lin, M., Juo, P., Hu, L., Anderson, M., Arden, K., Blenis, J. & Greenberg, M. (1999). Akt promotes cell survival by phosphorylating and inhibiting a Forkhead transcription factor. Cell, 96, 857–868.

Burgering, B. (2008). A brief introduction to FOXOlogy. Oncogene, 27, 2258–2262.

Calado, D., Sasaki, Y., Godinho, S., Pellerin, A., Köchert, K., Sleckman, B., Moreno de Alborán, I., Janz, M., Rodig, S. & Rajewsky, K. (2012). Th cell-cycle regulator c-Myc is essential for the formation and maintenance of germinal centers. Nat. Immunol. 13, 1092–1100.

Camacho, S. A., Kosco-Vilbois, M. H. & Berek, C. (1998). The Dynamic Structure of the Germinal Center. Immunol. Today, 19, 511–514.

de Boer, R. J. & Perelson, A. S. (2017). How Germinal Centers Evolve Broadly Neutralizing Antibodies: The Breadth of the Follicular Helper T Cell Response. J Virol, 91, e00983–17.

Dengler, H., Baracho, G., Omori, S., Bruckner, S., Arden, K., Castrillon, D., DePinho, R. & Rickert, R. (2008). Distinct functions for the transcription factor Foxo1 at various stages of B cell differentiation. Nat Immunol, 9, 1388–1398.

Depoil, D., Zaru, R., Guiraud, M., Chauveau, A., Harriague, J., Bismuth, G., Utzny, C., Muller, S. & Valitutti, S. (2005). Immunological synapses are versatile structures enabling selective T cell polarization. Immunity, 22, 185–194.

Dominguez-Sola, D., Kung, J., Holmes, A., Wells, V., Mo, T., Basso, K. & Dalla-Favera, R. (2015). The FOXO1 Transcription Factor Instructs the Germinal Center Dark Zone Program. Immunity, 43, 1064–1074.

Dominguez-Sola, D., Victora, G., Ying, C., Phan, R., Saito, M., Nussenzweig, M. & Dalla-Favera, R. (2012). The proto-oncogene MYC is required for selection in the germinal center and cyclic reentry. Nat Immunol, 13, 1083–1091.

Ersching, J., Efeyan, A., Mesin, L., Jacobsen, J., Pasqual, G., Grabiner, B., Dominguez-Sola, D., Sabatini, D. & Victora, G. (2017). Germinal Center Selection and Affinity Maturation Require Dynamic Regulation of mTORC1 Kinase. Immunity, 46, 1045–1058.

Figge, M. T., Garin, A., Gunzer, M., Kosco-Vilbois, M., Toellner, K.-M. & Meyer-Hermann, M. (2008). Deriving a germinal center lymphocyte migration model from two-photon data. J. Exp. Med. 205, 319–329.

Finkin, S., Hartweger, H., Oliveira, T. Y., Kara, E. E. & Nussenzweig, M. C. (2019). Protein Amounts of the MYC Transcription Factor Determine Germinal Center B Cell Division Capacity. Immunity, 51, 324–336.

Gitlin, A. D., Shulman, Z. & Nussenzweig, M. C. (2014). Clonal selection in the germinal centre by regulated proliferation and hypermutation. Nature, 509, 637–640.

Hannum, L. G., Haberman, A. M., Anderson, S. M. & Shlomchik, M. J. (2000). Germinal center initiation, variable gene region hypermutation, and mutant B cell selection without detectable immune complexes on follicular dendritic cells. J. Exp. Med. 192, 931–942.

Hauser, A. E., Junt, T., Mempel, T. R., Sneddon, M. W., Keinstein, S. H., Henrickson, S. E., von Andrian, U. H., Shlomchik, M. J. & Haberman, A. M. (2007). Definition of germinal-center B cell migration in vivo reveals predominant intrazonal circulation patterns. Immunity, 26, 655–667.

Heinzel, S., Binh Giang, T., Kan, A., Marchingo, J., Lye, B., Corcoran, L. & Hodgkin, P. (2017). A Myc-dependent division timer complements a cell-death timer to regulate T cell and B cell responses. Nat Immunol, 18, 96–103.

Hollmann, C. & Gerdes, J. (1999). Follicular Dendritic Cells and T-Cells - Nurses and Executioners in the Germinal Center Reaction. J. Pathol. 189, 147–149.

Hollowood, K. & Macartney, J. (1992). Cell kinetics of the germinal center reaction – a stathmokinetic study. Eur. J. Immunol. 22, 261–266.

Hur, D. Y., Kim, D. J., Kim, S., Kim, Y. I., Cho, D., Lee, D. S., Hwang, Y., Bae, K., Chang, K. Y. & Lee, W. J. (2000). Role of follicular dendritic cells in the apoptosis of germinal center B cells. Immunol. Lett. 72, 107–111.

Inoue, T., Shinnakasu, R., Ise, W., Kawai, C., Egawa, T. & Kurosaki, T. (2017). The transcription factor Foxo1 controls germinal center B cell proliferation in response to T cell help. J. Exp. Med. 214, 1181–1198.

Kuraoka, M., Schmidt, A., Nojima, T., Feng, F., Watanabe, A., Kitamura, D., Harrison, S., Kepler, T. & Kelsoe, G. (2016). Complex antigens drive permissive clonal selection in germinal centres. Immunity, 44, 542–552.

Lindhout, E., Lakeman, A. & de Groot, C. (1995). Follicular dendritic cells inhibit apoptosis in human B lymphocytes by a rapid and irreversible blockade of preexisting endonuclease. J. Exp. Med. 181 (6), 1985–1995.

Liu, D., Xu, H., Shih, C., Wan, Z., Ma, X., Ma, W., Luo, D. & Qi, H. (2015). T-B-cell entanglement and ICOSL-driven feed-forward regulation of germinal centre reaction. Nature, 517, 214–218.

Liu, Y. J., Zhang, J., Lane, P. J., Chan, E. Y. & MacLennan, I. C. M. (1991). Sites of specific B cell activation in primary and secondary responses to T cell-dependent and T cell-independent antigens. Eur. J. Immunol. 21, 2951–2962.

Luo, W., Weisel, F. & Shlomchik, M. (2018). B Cell Receptor and CD40 Signaling Are Rewired for Synergistic Induction of the c-Myc Transcription Factor in Germinal Center B Cells. Immunity, 48, 313–326.

MacLennan, I. C. M. (1994). Germinal Centers. Annu. Rev. Immunol. 12, 117–139.

Meyer-Hermann, M. (2014). Overcoming the dichotomy of quantity and quality in antibody responses. J. Immunol. 193, 5414–5419.

Meyer-Hermann, M. (2019). Injection of Antibodies against Immunodominant Epitopes Tunes Germinal Centers to Generate Broadly Neutralizing Antibodies. Cell Rep. 29, 1066–1073.

Meyer-Hermann, M., Binder, S., Mesin, L. & Victora, G. (2018). Computer simulation of multi-color Brainbow staining and clonal evolution of B cells in germinal centers. Front Immunol, 9, 2020.

Meyer-Hermann, M., Maini, P. K. & Iber, D. (2006). An analysis of B cell selection mechanisms in germinal centers. Math. Med. Biol. 23, 255–277.

Meyer-Hermann, M., Mohr, E., Pelletier, N., Zhang, Y., Victora, G. D. & Toellner, K.-M. (2012). A theory of germinal center B cell selection, division, and exit. Cell Rep. 2, 162–174.

Oprea, M., Nimwegen, E. v. & Perelson, A. S. (2000). Dynamics of One-pass Germinal Center Models: Implications for Affinity Maturation. Bull. Math. Biol. 62, 121–153.

Oprea, M. & Perelson, A. S. (1997). Somatic mutation leads to efficient affinity maturation when centrocytes recycle back to centroblasts. J. Immunol. 158, 5155–5162.

Sander, S., Chu, V., Yasuda, T., Franklin, A., Graf, R., Calado, D., Li, S., Imami, K., Selbach, M., Di Virgilio, M., Bullinger, L. & Rajewsky, K. (2015). PI3 Kinase and FOXO1 Tran-scription Factor Activity Differentially Control B Cells in the Germinal Center Light and Dark Zones. Immunity, 43, 1075–1086.

Schwickert, T. A., Lindquist, R. L., Schakhar, G., Livshits, G., Skokos, D., Kosco-Vilbois, M. H., Dustin, M. L. & Nussenzweig, M. C. (2007). In vivo imaging of germinal centres reveals a dynamic open structure. Nature, 446, 83–87.

Silver, J., Zuo, T., Chaudhary, N., Kumari, R., Tong, P., Giguere, S., Granato, A., Donthula, R., Devereaux, C. & Wesemann, D. (2018). Stochasticity enables BCR-independent germinal center initiation and antibody affinity maturation. J Exp Med, 215, 77–90.

Tas, J., Mesin, L., Pasqual, G., Targ, S., Jacobsen, J., Mano, Y., Chen, C., Weill, J., Reynaud, C., Browne, E., Meyer-Hermann, M. & Victora, G. (2016). Visualizing antibody affinity maturation in germinal centres. Science, 351, 1048–1054.

Tew, J. G., Wu, J., Qin, D., Helm, S., Burton, G. F. & Szakal, A. K. (1997). Follicular dendritic cells and presentation of antigen and costimulatory signals to B cells. Immunol Rev, 156, 39–52.

Turner, J., Ke, F. & Grigorova, I. (2018). B Cell Receptor Crosslinking Augments Germinal Center B Cell Selection when T Cell Help Is Limiting. Cell Rep, 25, 1395–1403.

Tzivion, G., Dobson, M. & Ramakrishnan, G. (2011). FoxO transcription factors; Regulation by AKT and 14-3-3 proteins. Biochim Biophys Acta, 1813, 1938–1945.

van Eijk, M., Defrance, T., Hennino, A. & de Groot, C. (2001). Death-receptor contribution to the germinal-center reaction. Trends Immunol. 22 (12), 677–682.

Victora, G., Dominguez-Sola, D., Holmes, A., Deroubaix, S., Dalla-Favera, R. & Nussenzweig, M. (2012). Identification of human germinal center light and dark zone cells and their relationship to human B-cell lymphomas. Blood, 120, 2240–2248.

Victora, G. D., Schwickert, T. A., Fooksman, D. R., Kamphorst, A. O., Meyer-Hermann, M., Dustin, M. L. & Nussenzweig, M. C. (2010). Germinal center dynamics revealed by multiphoton microscopy with a photoactivatable fluorescent reporter. Cell, 143, 592–605.

Vora, K., Ravetch, J. & Manser, T. (1997). Amplified follicular immune complex deposition in mice lacking the Fc receptor γ-chain does not alter maturation of the B cell response. J. Immunol. 159, 2116–2124.

Wang, P., Chang-ming, S., Qi, H. & Lan, Y.-h. (2016). A stochastic model of the germinal center integrating local antigen competition, individualistic T-B interactions, and B cell receptor signaling. J Immunol, 197, 1169–1182.

Wilhelm, K., Happel, K., Eelen, G., Schoors, S., Oellerich, M., Lim, R., Zimmermann, B., Aspalter, I., Franco, C., Boettger, T. & Fruttiger, M. e. a. (2016). FOXO1 couples metabolic activity and growth state in the vascular endothelium. Nature, 529, 216–220.

Zhang, J. & Shakhnovich, E. (2010). Optimality of Mutation and Selection in Germinal Centers. PLoS Comput Biol, 6, e1000800.

Zhang, Y., Meyer-Hermann, M., George, L., Khan, M., Figge, M. T., Falciani, F., Kosco-Vilbois, M. & Toellner, K.-M. (2013). Germinal centre B cells govern their own fate via antibody feedback. J. Exp. Med. 210, 457–464.

## References

Berek, C. & Milstein, C. (1987). Mutation drift and repertoire shift in the maturation of the immune response. Immunol. Rev. 96, 23–41.

Chaturvedi, A., Martz, R., Dorward, D., Waisberg, M. & Pierce, S. K. (2011). Endocytosed BCRs sequentially regulate MAPK and Akt signaling pathways from intracellular compartments. Nat. Immunol. 12, 1119–1126.

Dustin, M. & Meyer-Hermann, M. (2012). Antigen feast or famine. Science, 335, 408–409.

Meyer-Hermann, M. (2019). Injection of Antibodies against Immun-odominant Epitopes Tunes Germinal Centers to Generate Broadly Neutralizing Antibodies. Cell Rep. 29, 1066–1073.

Meyer-Hermann, M., Deutsch, A. & Or-Guil, M. (2001). Recycling Probability and Dynamical Properties of Germinal Center Reactions. J. Theor. Biol. 210, 265–285.

Miller, M., Wei, S., Parker, I. & Cahalan, M. (2002). Two-photon imaging of lymphocyte motility and antigen response in intact lymph node. Science, 296, 1869–73.

Nossal, G. (1992). The molecular and cellular basis of affinity maturation in the antibody response. Cell, 68, 1–2.

Omori, S. A., Cato, M. H., Anzelon-Mills, A., Puri, K. D., Shapiro-Shelef, M., Calame, K. & Richert, R. C. (2006). Regulation of Class-Switch Recombination and Plasma Cell Differentiation by Phosphatidylinositol 3-Kinase Signaling. Immunity, 25, 545–557.

Papa, I., Saliba, D., Ponzoni, M. & Bustamante, S. e. a. (2017). Tfh-derived dopamine accelerates productive synapses in germinal centres. Nature, 547, 318–323.

Perelson, A. S. & Oster, G. F. (1979). Theoretical Studies of Clonal Selection: Minimal Antibody Repertoire Size and Reliability of Self-Non-self Discrimination. J. Theor. Biol. 81, 645–670.

Thaunat, O., Granja, A. G., Barral, P., Filby, A., Montaner, B., Collinson, L., Nartinez-Martin, N., Harwood, N. E., Bruckbauer, A. & Batista, F. D. (2012). The asymmetric segregation of polarized antigen on B cell division shapes presentation capacity. Science, 335, 475–479.

Toellner, K. M., Jenkinson, W. E., Taylor, D. R., Khan, M., Sze, D. M. Y., Sansom, D. M., Vinuesa, C. G. & MacLennan, I. C. M. (2002). Low-level Hypermutation in T Cell-independent Germinal Centers Compared with High Mutation Rates Associated with T Cell-dependent Germinal Centers. J. Exp. Med. 195, 383–389.

